# Circulating miR-1285-3p promotes age-associated B cell differentiation through the OXPHOS-IKZF2 axis in SLE

**DOI:** 10.64898/2026.05.14.725263

**Authors:** Satoshi Akao, Hiromitsu Asashima, Hajime Inokuchi, Tatsuki Abe, Mohd Moin Khan, Nana Uematsu, Haruka Miki, Taihei Nishiyama, Ayako Ohyama, Yuya Kondo, Hiroto Tsuboi, Mineto Ota, Eliisa Kekäläinen, Kazuyoshi Ishigaki, Keishi Fujio, Isao Matsumoto

**Author notes:** **Corresponding author:** Hiromitsu Asashima, MD, PhD, Department of Rheumatology, Institute of Medicine, University of Tsukuba, 1-1-1 Tennodai, Tsukuba, Ibaraki 305-8575, Japan.

## Abstract

Age-associated B cells (ABCs) expand in systemic lupus erythematosus (SLE) and contribute to pathogenic humoral immunity, but the mechanisms that restrain their differentiation remain unclear. Here, we identify the transcription factor IKZF2 (Helios) as a regulator that limits ABC differentiation. Transcriptomic and functional analyses showed that suppression of oxidative phosphorylation (OXPHOS) in B cells promoted ABC differentiation and was accompanied by reduced *IKZF2* expression. Pharmacologic modulation of mitochondrial metabolism further demonstrated that OXPHOS inhibition promoted, whereas OXPHOS activation restrained, ABC differentiation. Integrative analyses revealed reduced *IKZF2* expression in selected B cell subsets from patients with SLE. Functional suppression of IKZF2 enhanced ABC differentiation and attenuated the inhibitory effects of OXPHOS activation, indicating that IKZF2 mediates metabolic control of B cell fate. Mechanistically, IKZF2 restrained early ABC-associated gene programs, including *ITGAX* and *TBX21*. Circulating miR-1285-3p in small extracellular vesicles, elevated in SLE, suppressed OXPHOS and recapitulated these effects. Together, these findings identify an OXPHOS-IKZF2 axis that restrains pathogenic B cell differentiation and links extracellular microRNA-mediated metabolic stress to ABC formation in SLE.

**One-sentence summary:** Small EV-associated miR-1285-3p in SLE promotes ABC differentiation by suppressing OXPHOS and relieving IKZF2-mediated restraint.

## Introduction

Systemic lupus erythematosus (SLE) is a heterogeneous autoimmune disease that predominantly affects women of childbearing age and involves multiple organs, including the skin, joints, and kidneys (*1*). A hallmark of SLE is the breakdown of B cell tolerance, which leads to the production of autoantibodies and the formation of immune complexes that drive systemic inflammation (*2*). Recent studies have demonstrated that aberrant B cell differentiation shapes the immunological landscape of SLE, underscoring the importance of B cell-intrinsic programs in disease development. Among B cell subsets, age-associated B cells (ABCs) have emerged as key drivers of SLE (*3*). In humans, ABCs are typically defined as IgD⁻CD27⁻ T-bet⁺ CD11c⁺ CD21⁻ B cells (*4*). The frequency of ABCs in the blood and affected tissues of patients with SLE has been shown to correlate with disease activity, including clinical manifestations and autoantibody production. ABC differentiation is primarily driven by Toll-like receptor 7 (TLR7) signaling, IFN-gamma (IFN-γ), and interleukin-21 (IL-21) (*5–8*). Furthermore, altered cellular metabolism, including defective mitochondrial remodeling in B cells, can promote an “aged” immune phenotype characterized by marked expansion of ABCs and a concomitant reduction in germinal center (GC) B cells in mice (*9*). Several transcription factors have been implicated in ABC function and differentiation. T-bet, a lineage-defining transcription factor, promotes the functional maturation of ABCs by driving immunoglobulin class switching in mouse and human B cells (*10, 11*). ZEB2, another key regulator, directly binds to the *ITGAX* (CD11c) promoter and contributes to the acquisition of the CD11c⁺ phenotype characteristic of ABCs (*12*). However, the upstream signals that govern ABC development remain incompletely understood.

Small extracellular vesicles (small EVs), typically less than 200 nm in diameter, have gained attention as potent mediators of intercellular communication. Circulating small EVs are increased in patients with SLE and correlate with disease activity, as measured by the SLE Disease Activity Index (SLEDAI) (*13*). Although small EV-derived microRNAs (miRNAs) are known to activate plasmacytoid dendritic cells and amplify type I IFN responses (*14*), their direct effects on B cell differentiation and metabolic programming remain poorly understood. Notably, small EV-derived miRNAs can modulate mitochondrial pathways, including oxidative phosphorylation (OXPHOS), suggesting that they may influence immune cell fate decisions (*15, 16*). However, the contribution of miRNAs to ABC differentiation in SLE has not been elucidated.

Here, we identify a mechanism by which circulating miR-1285-3p, elevated in small EVs from patients with SLE, promotes ABC differentiation. We show that miR-1285-3p represses OXPHOS-related metabolic pathways and downregulates the transcription factor IKZF2, leading to increased expression of CD11c and T-bet. Our findings reveal IKZF2 as a novel regulator of ABC formation and establish a metabolic-transcriptional axis linking miRNA signaling to pathogenic B cell differentiation in SLE.

## Results

### Identification of circulating small EV-derived miRNAs in SLE

To explore disease-associated miRNAs contained in circulating small EVs, we first isolated them from plasma samples of treatment-naïve patients with SLE and age- and sex-matched HCs (n = 4 per group) (Fig. 1A). Nanoparticle tracking analysis showed a particle size distribution consistent with small EVs (mean, 180 ± 6.8 nm; mode, 156 ± 5.9 nm) (Fig. 1B). Transmission electron microscopy confirmed round, membrane-bound vesicles less than 100 nm in diameter (Fig. 1C), and western blotting detected canonical EV markers CD9 and CD63, but not the intracellular protein cytochrome c (Fig. 1D). These findings validate the successful isolation of small EVs with minimal contamination.

**Fig. 1.**
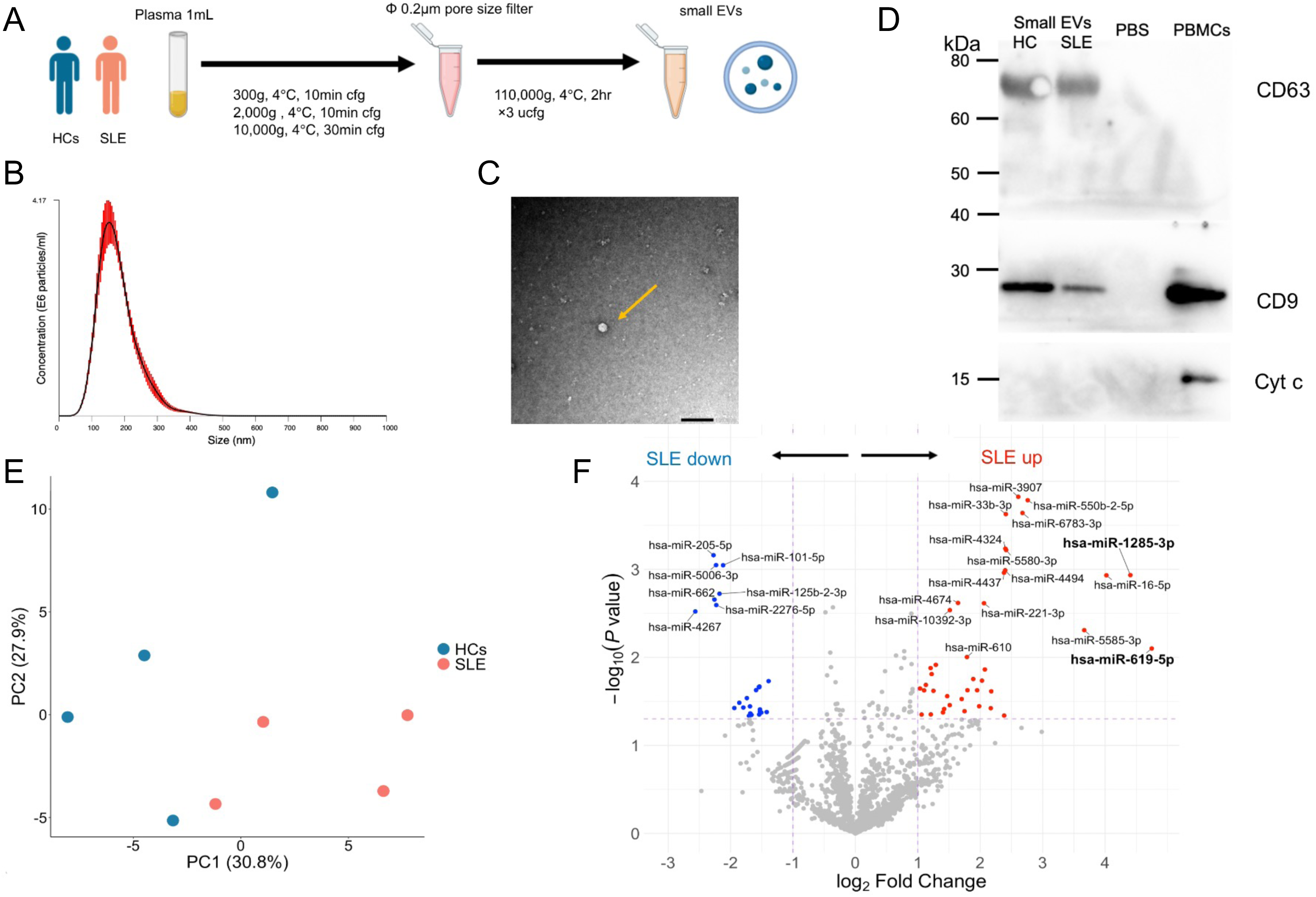
Characterization of plasma-derived small EVs and identification of miRNAs in SLE. (**A**) Workflow for isolation of plasma-derived small EVs from treatment-naïve patients with systemic lupus erythematosus (SLE) and age- and sex-matched healthy controls (HCs) (n = 4 per group). (**B**) Nanoparticle tracking analysis (NTA) of isolated EVs showing a particle size distribution consistent with small EVs (mean, 180 ± 6.8 nm; mode, 156 ± 5.9 nm). (**C**) Transmission electron microscopy (TEM) images demonstrating round, membrane-bound vesicles. The yellow arrow indicates a representative vesicle. Scale bar, 100 nm. (**D**) Western blot analysis confirming the presence of canonical EV markers CD9 and CD63 and the absence of the intracellular protein cytochrome c in SLE- and HC-derived small EVs. (**E**) Principal component analysis of microarray-based miRNA profiles exhibiting clear separation between SLE-derived (red) and HC-derived (blue) small EVs (n = 4 per group). (**F**) Volcano plot of differentially expressed EV miRNAs between patients with SLE and HCs. miRNAs downregulated in SLE (*P* < 0.05, log₂ fold change < −1) are shown in blue, upregulated miRNAs (*P* < 0.05, log₂ fold change > 1) in red, and all others in gray. miRNAs with −log₁₀(*P* value) > 2 are annotated, highlighting miR-1285-3p and miR-619-5p as the most significantly upregulated miRNAs in SLE-derived small EVs and selected for further validation. cfg, centrifuge; ucfg, ultracentrifuge; kDa, kilodalton; PBS, phosphate-buffered saline; PBMCs, peripheral blood mononuclear cells; Cyt c, cytochrome c.

We next performed microarray-based miRNA profiling using these small EVs. Principal component analysis (PCA) revealed distinct clustering between the two groups, indicating disease-associated alterations in miRNA composition (Fig. 1E). Differential expression analysis identified 40 upregulated and 25 downregulated miRNAs in SLE-derived small EVs compared with HCs. Among all differentially expressed miRNAs, miR-1285-3p and miR-619-5p were the most significantly upregulated in SLE-derived small EVs and were therefore selected for further validation (Fig. 1F).

### miR-1285-3p drives ABC differentiation and reflects disease activity in SLE

To further evaluate the impact of these miRNAs on B cell differentiation, naïve B cells were transfected with either miRNA mimics (miR-1285-3p or miR-619-5p) or a negative control and cultured under conditions that promote ABC differentiation (Fig. 2A and Supplementary Fig. 1). Whereas miR-619-5p transfection did not alter the frequency of ABCs, miR-1285-3p transfection resulted in a significant increase in the proportion of ABCs (Fig. 2, B and C). Details of transfection efficiency are provided in the Supplementary Figure 2.

**Fig. 2.**
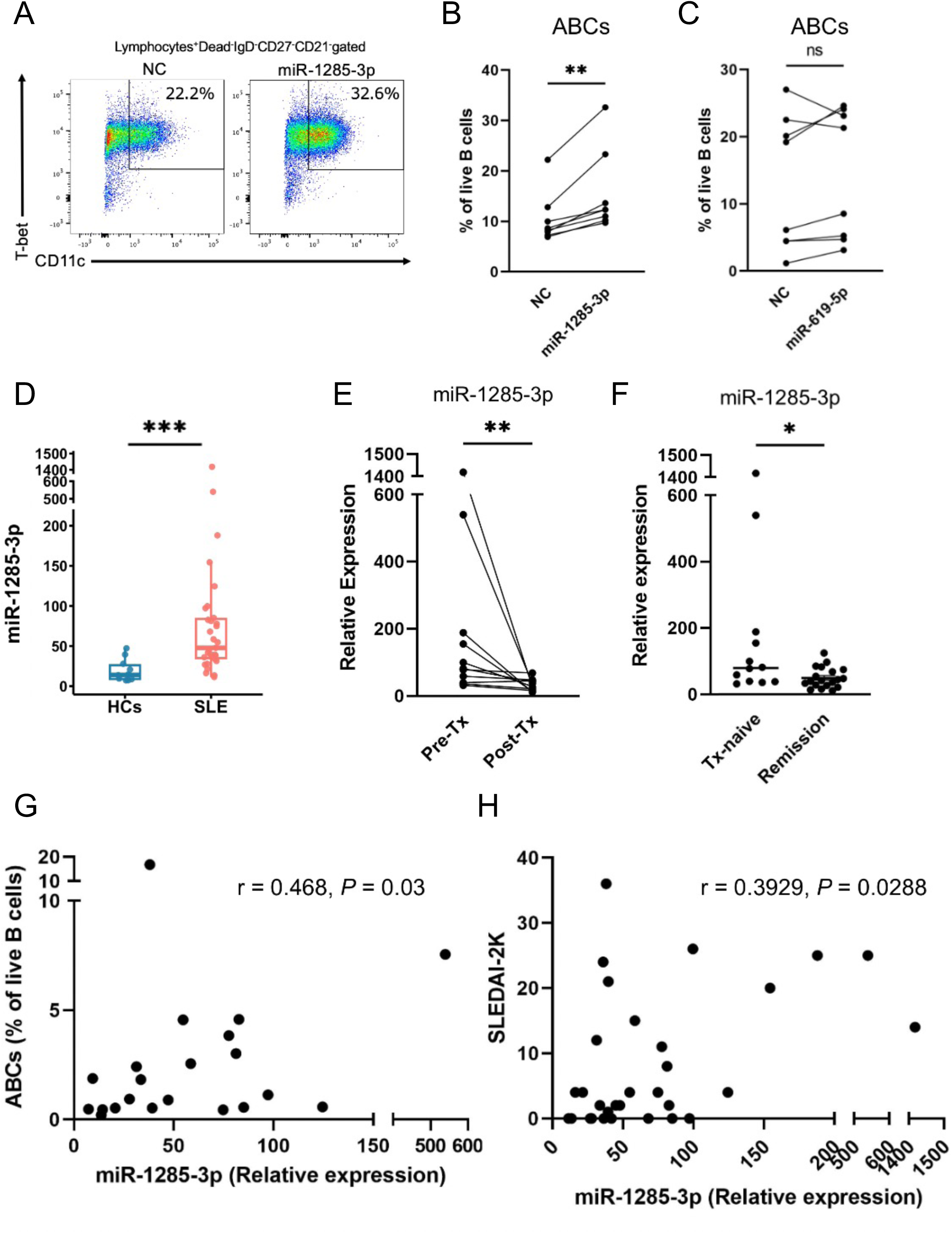
miR-1285-3p promotes ABC differentiation *in vitro* and associates with immunological and clinical activity in SLE. (**A**) Representative flow cytometry plots showing frequencies of ABCs (CD11c⁺T-bet⁺) after miR-1285-3p transfection. (**B** and **C**) Frequency of ABCs among live B cells under ABC-inducing culture conditions after transfection with miR-1285-3p mimic or negative control (NC). Each line represents paired samples from an independent experiment using HC-derived naïve B cells (n = 8). (**D**) Expression of miR-1285-3p in plasma-derived small EVs from treatment-naïve patients with SLE (n = 31) and HCs (n = 10). (**E**) Longitudinal analysis of miR-1285-3p expression in plasma-derived small EVs before and after treatment in treatment-naïve patients with SLE (n = 11). (**F**) Comparison of miR-1285-3p expression between patients with active SLE (n = 12) and patients who achieved Lupus Low Disease Activity State (LLDAS; n = 19). (**G**) Relationship between the frequency of circulating ABCs and miR-1285-3p expression in plasma-derived small EVs in patients with SLE (n = 22). (**H**) Relationship between SLEDAI-2K scores and miR-1285-3p expression in plasma-derived small EVs (n = 31). Data are presented as mean ± SD. \**P* < 0.05, \*\**P* < 0.01, \*\*\**P* < 0.001; ns, not significant. Statistical analysis was performed using the Wilcoxon test for (**B**), (**C**), and (**E**); the Mann–Whitney U test for (**D**) and (**F**); and Spearman correlation for (**G**) and (**H**). ABCs, age-associated B cells; NC, negative control; Tx, treatment; SLEDAI-2K, Systemic Lupus Erythematosus Disease Activity Index 2000.

Because previous studies have shown that ABCs are expanded in patients with SLE (*6, 17, 18*) and only miR-1285-3p enhanced ABC differentiation, we next evaluated its clinical relevance in SLE. As shown in Figure 1F, small EVs from patients with SLE exhibited significantly higher levels of miR-1285-3p than those from HCs (Fig. 2D). In treatment-naïve patients with SLE, expression of miR-1285-3p in plasma-derived small EVs significantly decreased after treatment (Fig. 2E). We also observed a significant difference in miR-1285-3p expression between patients with active SLE and those who had achieved the Lupus Low Disease Activity State (LLDAS) (Fig. 2F). Moreover, miR-1285-3p expression showed a significant positive correlation with both the frequency of circulating ABCs and the SLEDAI-2K score (Fig. 2, G and H).

Together, these findings suggest that miR-1285-3p promotes ABC differentiation *in vitro* and is elevated in plasma-derived small EVs from patients with SLE, where it is associated with immunological and clinical disease activity.

### OXPHOS suppression underlies miR-1285-3p-mediated induction of ABCs

We next examined the mechanism by which miR-1285-3p promotes ABC differentiation. Naïve B cells transfected with miR-1285-3p or a negative control and cultured under ABC-inducing conditions were harvested for bulk RNA-seq analysis (n = 5 per group). Because miRNAs generally suppress the expression of their target messenger RNAs (mRNAs), pathway analysis focused on genes downregulated in the miR-1285-3p group (*P* < 0.1, log₂FC < −0.5). This analysis identified OXPHOS signaling as a significantly enriched pathway among downregulated genes (Fig. 3A). A volcano plot confirmed broad suppression of OXPHOS-related genes in miR-1285-3p-transfected cells (Fig. 3B). Consistent with these findings, pathway analysis of downregulated genes from naïve B cells of patients with SLE (GSE92387) also revealed significant enrichment of OXPHOS-related genes (Supplementary Fig. 3), indicating that OXPHOS suppression represents a shared transcriptional feature between miR-1285-3p-transfected B cells and naïve B cells in SLE.

**Fig. 3.**
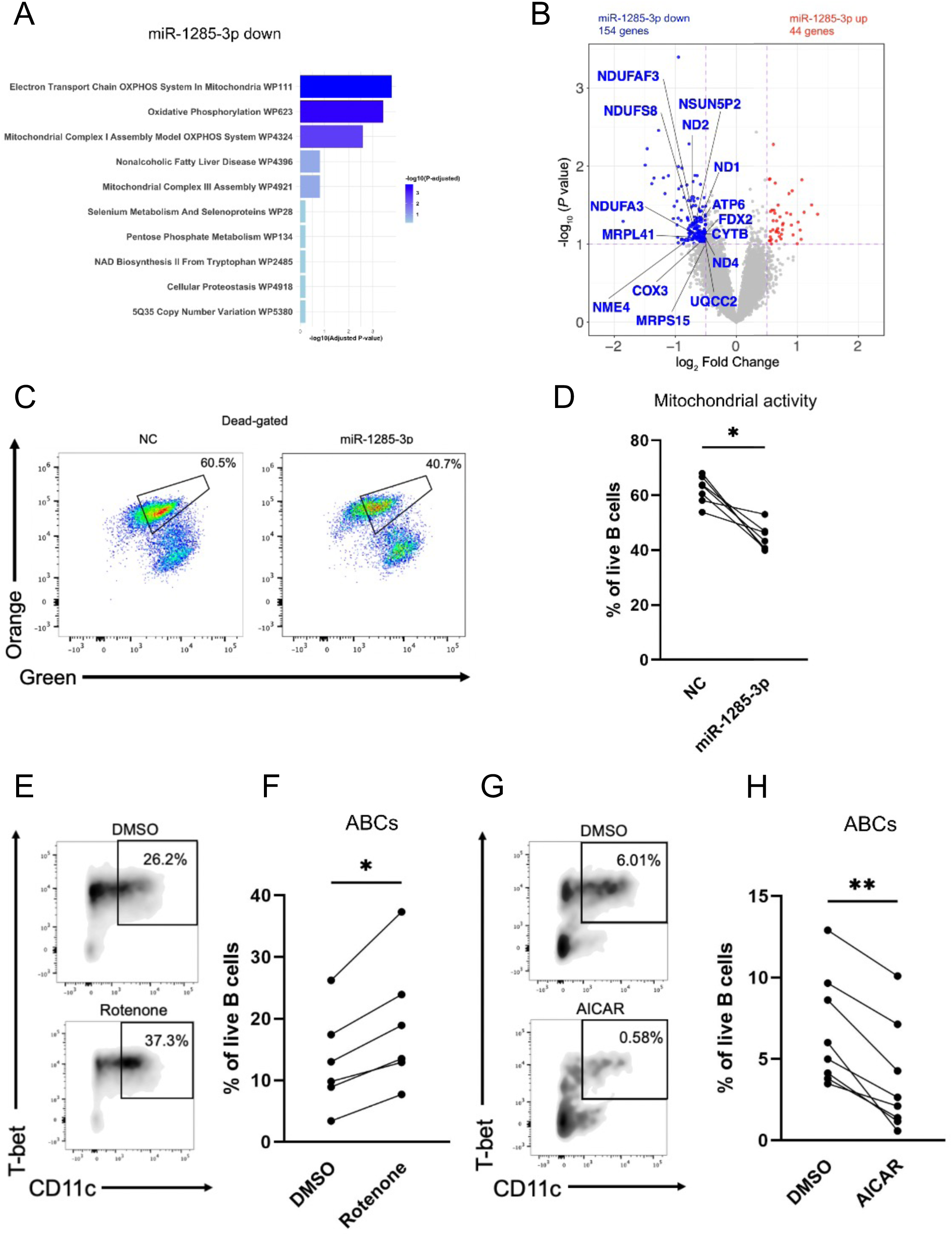
miR-1285-3p promotes ABC differentiation through suppression of mitochondrial OXPHOS signaling. (**A**) Pathway enrichment analysis of genes downregulated in miR-1285-3p-transfected B cells compared with negative control under ABC-inducing conditions (n = 5 per group). Analysis focused on genes with *P* < 0.1 and log₂ fold change < −0.5. Pathways are ranked by −log₁₀(*P* value). (**B**) Volcano plot of differentially expressed genes from bulk RNA-seq analysis. Genes downregulated in the miR-1285-3p group (*P* < 0.1, log₂ fold change < −0.5) are shown in blue, upregulated genes (*P* < 0.1, log₂ fold change > 0.5) in red, and all others in gray. All annotated genes represent OXPHOS-related genes. (**C**) Representative flow cytometry plots of mitochondrial mass and membrane potential assessed by MitoSpy Green and MitoSpy Orange staining in live B cells transfected with miR-1285-3p or negative control. The boxed population denotes cells with enhanced mitochondrial mass and membrane potential, indicating elevated mitochondrial activity. (**D**) Quantification of the proportion of cells with activated mitochondrial function, showing a significant reduction in the miR-1285-3p-transfected group (n = 6 per group). (**E**) Representative flow cytometry plots showing frequencies of ABCs (CD11c⁺T-bet⁺) following inhibition of mitochondrial complex I with rotenone or vehicle control (DMSO). (**F**) Quantification of ABC frequencies in (**E**) (n = 6 per group). (**G**) Representative flow cytometry plots showing frequencies of ABCs following activation of OXPHOS with AICAR or DMSO. (**H**) Quantification of ABC frequencies in (**G**) (n = 8 per group). Data are presented as mean ± SD. \**P* < 0.05, \*\**P* < 0.01; ns, not significant. Wilcoxon signed-rank test was used for (**D**), (**F**), and (**H**).

To functionally assess mitochondrial metabolism, mitochondrial mass and membrane potential were evaluated using MitoSpy Green and MitoSpy Orange, established indicators of mitochondrial activity, in activated B cells (*19*). The proportion of cells with activated mitochondrial function was reduced in the miR-1285-3p group (Fig. 3, C and D). Modulation of mitochondrial metabolism further supported a role for OXPHOS in ABC differentiation. Inhibition of complex I with rotenone increased the frequencies of ABCs (Fig. 3, E and F), whereas OXPHOS activation with AICAR suppressed ABC induction (Fig. 3, G and H). In contrast, ATP synthase inhibition with oligomycin had no appreciable effect on ABC differentiation (Supplementary Fig. 4), suggesting that ATP depletion alone is insufficient to drive this process and that disrupted electron flow at complex I provides a more proximal signal for ABC differentiation.

To further examine whether mitochondrial perturbation could directly promote ABC-associated features in B cells, we used the human B cell line Raji. Although Raji cells do not fully recapitulate ABC differentiation, rotenone treatment induced a concordant change in CD11c expression, consistent with the findings in primary B cells (Supplementary Fig. 5A). In addition, ABC-inducing conditions also enhanced CD11c expression, which was further augmented by the addition of rotenone (Supplementary Fig. 5B). Thus, an independent B cell line model recapitulated the core effect of mitochondrial perturbation observed in primary naïve B cells, supporting the reproducibility of this phenotype. These findings suggest that mitochondrial dysfunction is sufficient to drive key ABC-associated phenotypic changes in a B cell-intrinsic manner.

Because complex I inhibition can trigger ROS generation, we assessed whether acute mtROS generation accompanies complex I inhibition. MitoSOX staining performed 1, 2, and 4 hours after rotenone or oligomycin treatment revealed no substantial increase in mtROS levels compared with control (Supplementary Fig. 6A), indicating that early mtROS surges are unlikely to account for the observed ABC expansion. However, coadministration of the ROS scavenger N-acetylcysteine (NAC) partially attenuated rotenone-induced ABC differentiation (Supplementary Fig. 6B), suggesting that although acute mtROS spikes were not detected, ROS-related signaling downstream of mitochondrial complex I inhibition may still contribute to ABC induction.

### OXPHOS inhibition downregulates IKZF2 and relieves a brake on ABC differentiation

To investigate the mechanism by which OXPHOS inhibition promotes ABC differentiation, bulk RNA-seq was performed on cells harvested 24 hours after rotenone treatment under ABC-inducing conditions. To compare these transcriptional changes with those observed in SLE, we analyzed transcriptomic data from naïve B cells in the ImmuNexUT cohort, which contains sorted B cell subsets from patients with SLE and HCs (JGAS000486) (*20*). This integrative analysis identified seven transcription factors commonly upregulated (*BHLHE41*, *FOS*, *HIVEP1*, *IRF1*, *KLF6*, *KLF10*, and *TSC22D1*) and one downregulated (*IKZF2*) across both datasets (Fig. 4A). These findings highlight IKZF2 as a candidate regulator linking metabolic stress to ABC differentiation in SLE.

**Fig. 4.**
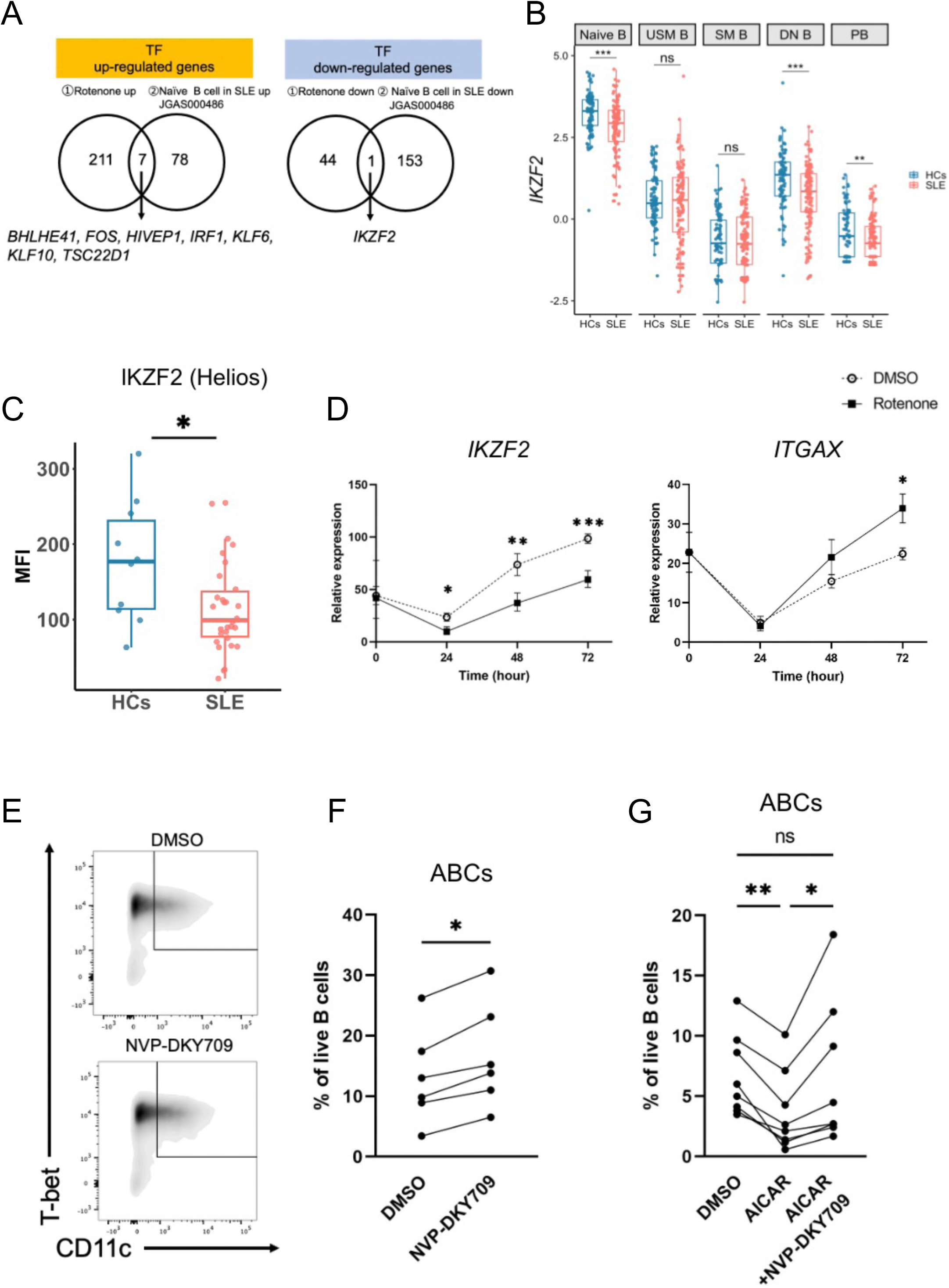
OXPHOS inhibition promotes ABC differentiation through downregulation of IKZF2. (**A**) Integrative transcription factor (TF) analysis comparing bulk RNA-seq data from B cells treated with rotenone under ABC-inducing conditions (n = 4 per group) and publicly available RNA-seq data from naïve B cells of patients with SLE and healthy controls (HCs) in the ImmuNexUT cohort (JGA S000486). Venn diagrams show TFs commonly upregulated (left) or downregulated (right) in both datasets (*P* < 0.01). (**B**) *IKZF2* mRNA expression in naïve, unswitched memory (USM), switched memory (SM), double-negative (DN) B cells, and plasmablasts (PBs) from patients with SLE and healthy controls (HCs) in the ImmuNexUT cohort. Sample numbers for each subset were as follows: naïve B cells (SLE n = 128, HCs n = 88), USM B cells (SLE n = 128, HCs n = 89), SM B cells (SLE n = 135, HCs n = 89), DN B cells (SLE n = 134, HCs n = 87), and PBs (SLE n = 130, HCs n = 88). (**C**) Flow cytometric analysis of IKZF2 (Helios) expression in ABCs from patients with SLE (n = 30) and HCs (n = 10). (**D**) Time-course analysis of ABC-associated gene expression following OXPHOS inhibition with rotenone (n = 6 per group). (**E**) Representative flow cytometry plots showing frequencies of ABCs (CD11c⁺T-bet⁺) following treatment with the selective IKZF2 molecular glue degrader NVP-DKY709 or DMSO. (**F**) Quantification of ABC frequencies in (**E**) (n = 6 per group). (**G**) Quantification of ABC frequencies in B cells treated with DMSO, AICAR, or AICAR in combination with NVP-DKY709 (n = 8 per group). Data are presented as mean ± SD. \**P* < 0.05, \*\**P* < 0.01, \*\*\**P* < 0.001; ns, not significant. Mann–Whitney U test was used for (**C**); two-way ANOVA with multiple comparisons for (**D**); and Wilcoxon signed-rank test for (**F**) and (**G**). ABCs, age-associated B cells; HCs, healthy controls.

*IKZF2*, which encodes Helios, has been reported as an SLE susceptibility gene and is implicated in immune regulation through Helios⁺ Treg cells (*21, 22*). Although less well characterized in B cells, *IKZF2* is expressed in the B cell lineage, and loss-of-function mutations have been associated with autoimmunity (*23*). These observations led us to assess *IKZF2* expression following OXPHOS inhibition. qPCR confirmed a significant reduction in *IKZF2* expression at 24 hours after rotenone treatment compared with DMSO-treated controls (Supplementary Fig. 7).

To further evaluate the clinical relevance of IKZF2, we next performed a complementary analysis of IKZF2 expression across B cell subsets in the ImmuNexUT cohort. *IKZF2* expression was significantly lower in naïve B cells, double-negative (DN) B cells, and plasmablasts from patients with SLE than in healthy controls, whereas no significant differences were observed in other B cell subsets (Fig. 4B). These findings indicate that *IKZF2* downregulation is not uniform across B cell compartments in SLE but is most evident in selected subsets, particularly naïve and DN B cells, that are linked to disease-associated ABC differentiation.

Flow cytometric analysis revealed that IKZF2 (Helios) expression was also significantly decreased in ABCs from patients with SLE compared with healthy controls (Fig. 4C), supporting the transcriptional findings and suggesting reduced *IKZF2* expression in pathogenic B cell states.

We next examined the temporal dynamics of ABC-associated genes after OXPHOS inhibition. *IKZF2* expression remained suppressed from 24 to 72 hours, whereas *ITGAX* (CD11c) expression increased beginning at 48 hours (Fig. 4D). *TBX21* (T-bet) exhibited early downregulation at 24 and 48 hours followed by recovery at 72 hours and further upregulation at 96 hours, consistent with delayed induction during ABC differentiation (Supplementary Fig. 8). No significant changes were detected in *ZEB2* expression (Supplementary Fig. 9).

To functionally test the role of IKZF2, we perturbed its expression through pharmacologic and genetic approaches. Treatment with NVP-DKY709, a selective IKZF2 molecular glue degrader, increased the proportion of ABCs (Fig. 4, E and F). Consistently, IKZF2 knockdown by siRNA similarly enhanced ABC differentiation (Supplementary Fig. 10, A and B). Notably, coadministration of NVP-DKY709 attenuated the inhibitory effect of AICAR on ABC differentiation (Fig. 4G), indicating that IKZF2 mediates the effects of OXPHOS suppression on ABC differentiation.

Together, these findings demonstrate that OXPHOS inhibition reduces *IKZF2* expression and that diminished IKZF2 activity facilitates ABC differentiation.

### IKZF2 restrains early ABC-associated gene programs in SLE

We next examined how IKZF2 constrains the expression of ABC-related genes. Following IKZF2 degradation with NVP-DKY709, we quantified the expression of ABC-associated transcripts over time. *ITGAX* and *TBX21* showed little change at 24 and 48 hours but were robustly induced at 72 hours, whereas *ZEB2* expression remained unchanged throughout the time course (Fig. 5, A and B, and Supplementary Fig. 11A).

**Fig. 5.**
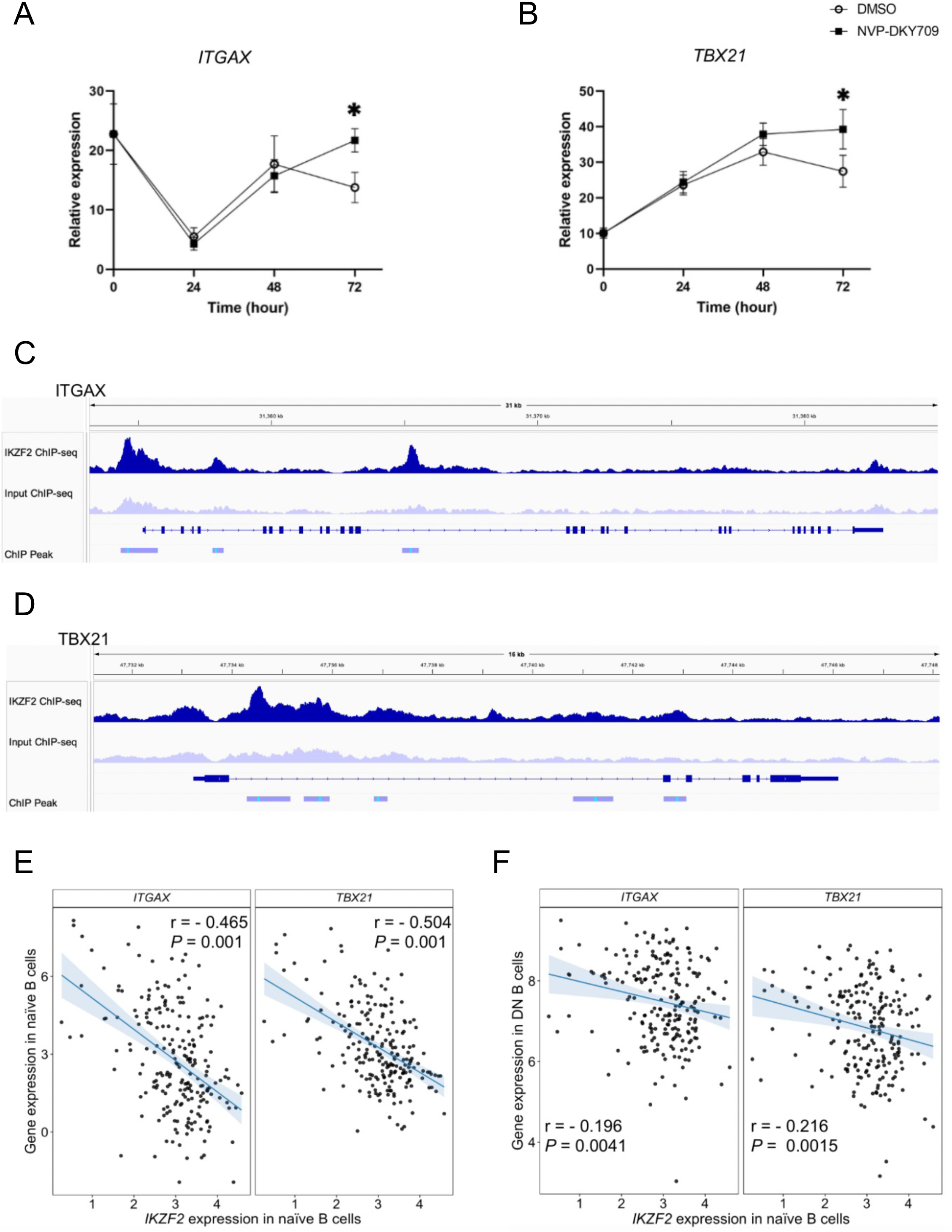
IKZF2 regulates ABC-associated genes *ITGAX* and *TBX21*. (**A** and **B**) Time-course qPCR analysis of *ITGAX* (**A**) or *TBX21* (**B**) expression in B cells treated with NVP-DKY709 or DMSO under ABC-inducing conditions (n = 6 per group). (**C** and **D**) Representative genome browser tracks showing IKZF2 ChIP-seq signal, input control (ENCSR822AHX), and called peaks across the *ITGAX* (**C**) or *TBX21* (**D**) loci in the human B cell line GM12878. (**E**) Correlation between *IKZF2* expression in naïve B cells and *ITGAX* or *TBX21* expression within the same subset from patients with SLE. (**F**) Correlation between *IKZF2* expression in naïve B cells and *ITGAX* or *TBX21* expression in double-negative (DN) B cells from patients with SLE. Each dot represents an individual sample. Sample numbers for each subset were as follows: naïve B cells (n = 128) and DN B cells (n = 134). Data are presented as mean ± SD. \**P* < 0.05; ns, not significant. Mann-Whitney U test was used for (**A**) and (**B**); Pearson correlation was used for (**E**) and (**F**).

Given that *ITGAX* and *TBX21* were selectively induced after IKZF2 degradation, we next assessed whether IKZF2 could directly associate with these loci. Reanalysis of public IKZF2 ChIP-seq data from a human B cell line (ENCSR822AHX) revealed prominent *IKZF2* occupancy at the *ITGAX* and *TBX21* loci, including peaks near promoter-associated regions, and also at the *ZEB2* locus (Fig. 5, C and D, and Supplementary Fig. 11B). These findings suggest that *IKZF2* can access regulatory regions of key ABC-associated genes, although transcriptional induction after IKZF2 degradation was observed only for *ITGAX* and *TBX21* under the conditions tested.

To determine whether IKZF2 is linked to ABC-associated genes in SLE, we analyzed correlations between *IKZF2* expression and ABC-associated genes in B cell subsets from patients with SLE using transcriptomic data from the ImmuNexUT cohort (*20*). *IKZF2* expression in naïve B cells showed a moderate inverse correlation with *ITGAX* and *TBX21* expression within the same subset (Fig. 5E). In addition, *IKZF2* expression in naïve B cells exhibited a weaker but consistent inverse correlation with *ITGAX* and *TBX21* expression in DN B cells (Fig. 5F). By contrast, *IKZF2* expression in DN B cells did not correlate with *ITGAX* or *TBX21* expression within the same subset (Supplementary Fig. 12).

These findings support a model in which IKZF2 restrains ABC-associated gene programs primarily at the naïve B cell stage, such that reduced *IKZF2* expression in naïve B cells in SLE may release this restraint and facilitate differentiation toward the ABC state.

### Germline IKZF2 loss-of-function mutation drives ABC expansion

To provide human genetic evidence supporting a causal role of IKZF2 in restraining ABC differentiation, we analyzed B cell phenotypes in individuals harboring a heterozygous IKZF2 loss-of-function mutation (p.Y200X), identified in a previously reported Finnish family cohort (*24*). The pedigree demonstrated the presence of the IKZF2 nonsense mutation across two generations within the affected family (Fig. 6A). Flow cytometric analysis revealed an elevated frequency of ABCs among CD19⁺ B cells in both individuals carrying the IKZF2 mutation compared with HCs (Fig. 6, B and C). The expansion of ABCs was accompanied by a broader redistribution of peripheral B cell subsets, with an increase of naïve B cells and reciprocal reductions in memory B cells in IKZF2 mutation carriers relative to HCs (Supplementary Fig. 13).

**Fig. 6.**
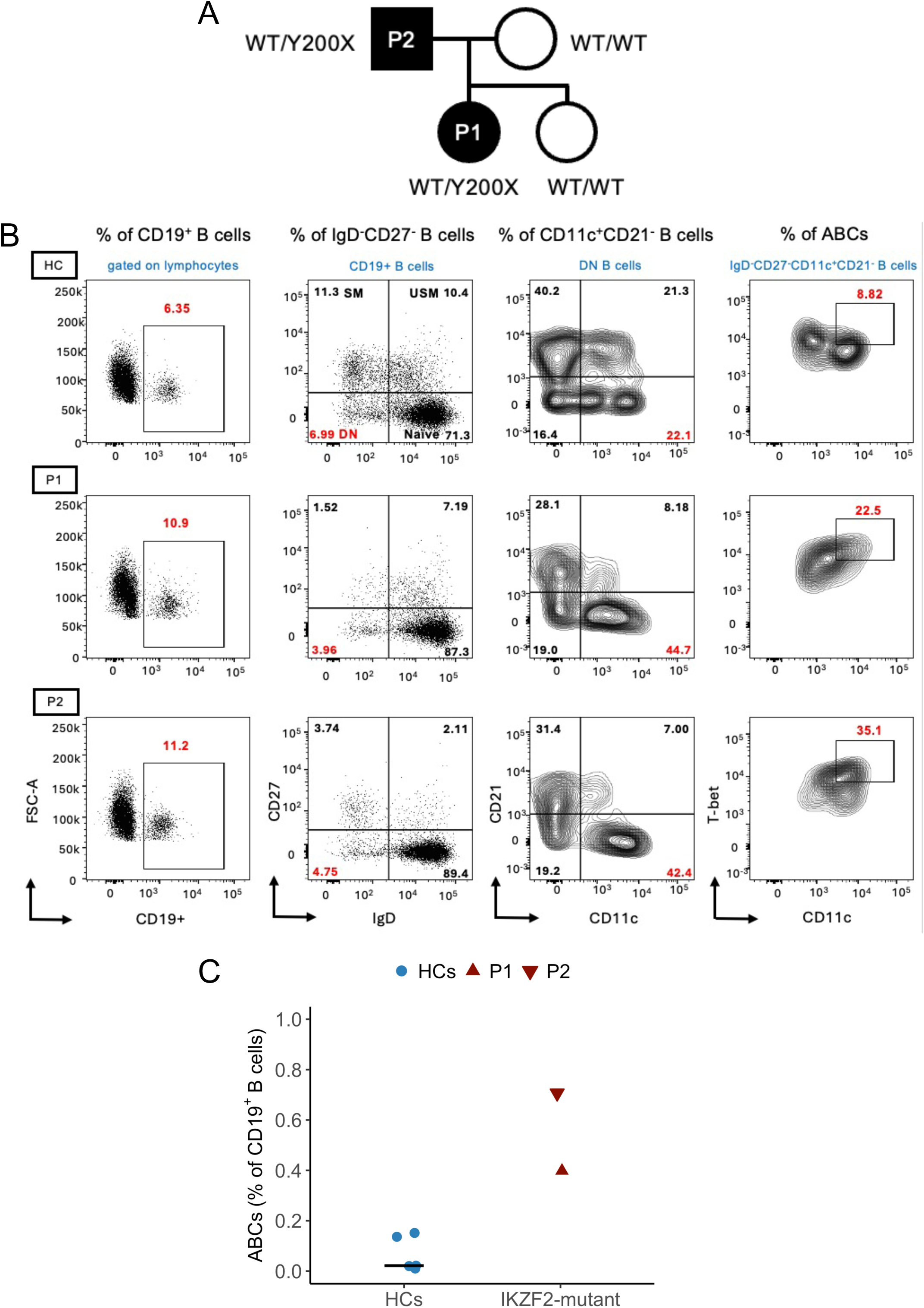
Increased frequency of ABCs in individuals carrying an IKZF2 loss-of-function mutation. (**A**) Pedigree of a family carrying an IKZF2 loss-of-function mutation. The pedigree shows two generations. Squares represent males and circles represent females. The proband (P1) and her father (P2) are affected and carry the heterozygous IKZF2 nonsense mutation (c.600C>A; p.Y200X). (**B**) Representative flow cytometry plots illustrating the gating strategy for B cell analysis, including identification of CD19⁺ B cells, DN cells, and ABCs in individuals carrying the IKZF2 mutation (P1 and P2) and a healthy control (HC). (**C**) Frequencies of ABCs among CD19⁺ B cells in individuals carrying the IKZF2 mutation (P1 and P2) and HCs. Each dot represents an individual sample. Horizontal bars indicate the median. USM, Unswitched Memory B cells; SM, Switched Memory B cells; DN, double-negative B cells; ABCs, age-associated B cells; IKZF2-mutant, individuals carrying the IKZF2 mutation; HCs, healthy controls.

Together, these findings provide in vivo human genetic evidence supporting a role for IKZF2 in shaping B cell fate, with a shift toward ABC-like phenotypes.

## Discussion

In this study, we identified an OXPHOS–IKZF2 regulatory axis that restrains ABC-associated B cell differentiation in SLE, with small EV-associated miR-1285-3p acting as a potential upstream modulator. miR-1285-3p was selectively enriched in circulating small EVs from patients with SLE, correlated with clinical disease activity, and functionally enhanced ABC differentiation. Mechanistically, miR-1285-3p suppressed OXPHOS in B cells, reduced *IKZF2* expression, and thereby relieved restraint of early ABC-associated gene programs, including *ITGAX* and *TBX21*. Together, these findings support a model in which small EV-associated miRNAs shape pathogenic B cell fate through an OXPHOS-IKZF2 axis (Fig. 7).

**Fig. 7.**
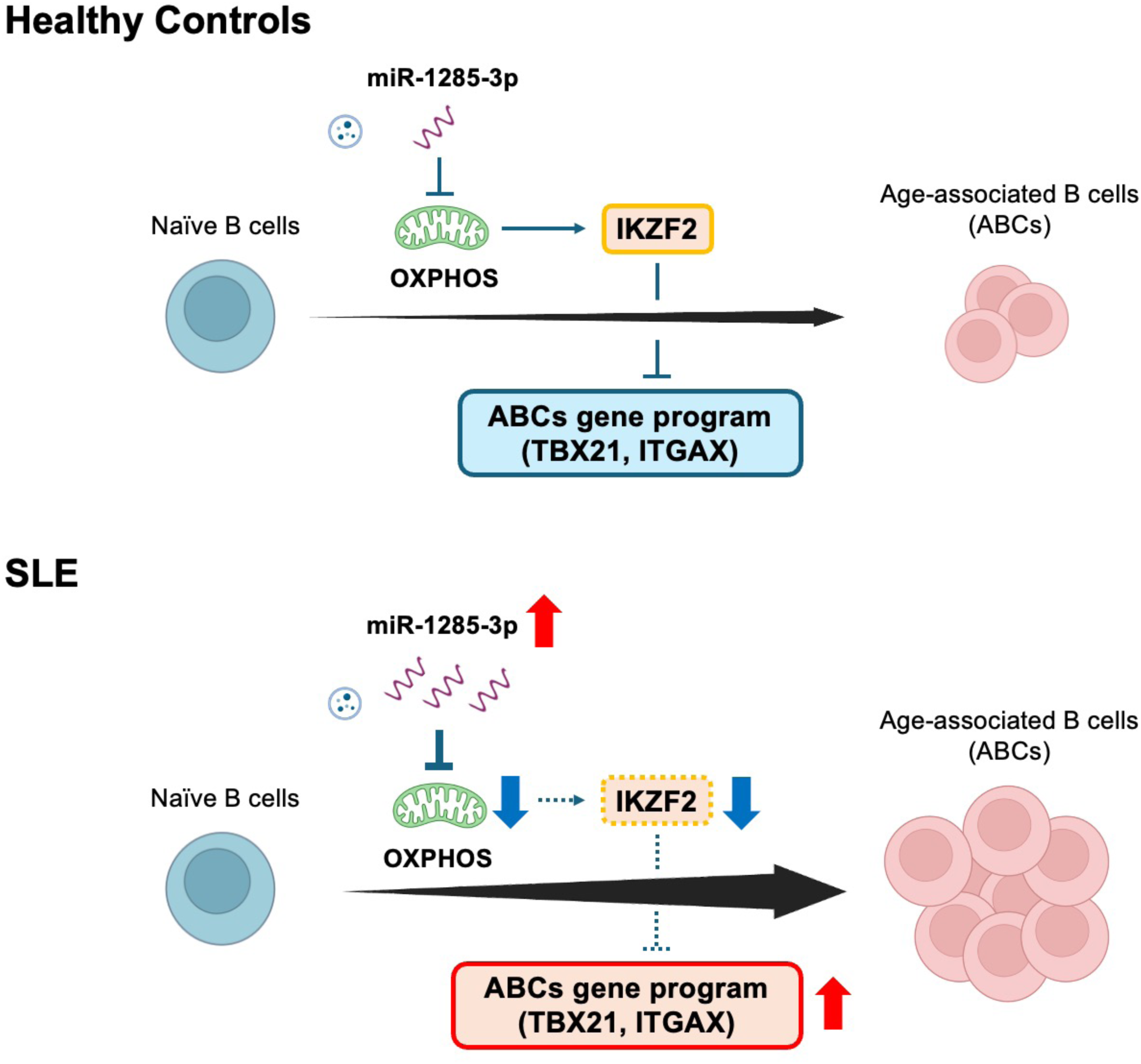
Graphical abstract. Schematic model illustrating how metabolic reprogramming regulates age-associated B cell (ABC) differentiation in healthy conditions and in SLE. In healthy B cells, mitochondrial oxidative phosphorylation (OXPHOS) is maintained, supporting expression of the transcription factor IKZF2 (Helios), which acts as a transcriptional brake to restrain ABC-associated genes, including *ITGAX* (CD11c) and *TBX21* (T-bet), thereby limiting ABC differentiation. In SLE, plasma-derived small extracellular vesicles (EVs) enriched in miR-1285-3p suppress OXPHOS, leading to downregulation of *IKZF2*. Loss of *IKZF2* releases transcriptional repression of *ITGAX* and *TBX21*, promoting ABC expansion. This model highlights a metabolic–IKZF2 axis linking mitochondrial dysfunction to pathogenic B cell differentiation in SLE.

Small EVs are increasingly recognized as mediators of intercellular communication (*25–27*). In SLE, small EVs have been implicated in immune activation through nucleic acids and innate sensing and can influence adaptive immune functions (*14, 28*). However, the extent to which small EV-associated miRNAs directly regulate B cell differentiation has remained unclear. Our data suggest that miR-1285-3p may act as one of the upstream signals linking extracellular vesicle biology to mitochondrial and transcriptional remodeling in B cells. Its enrichment in small EVs is consistent with regulated packaging and raises the possibility that inflammatory cues in SLE promote the export of miR-1285-3p into the circulation. Defining the cellular sources of these vesicles and the upstream signals that control miR-1285-3p loading will be an important next step.

Immunometabolism can instruct cell fate and gene expression (*29*). Prior work has shown that ABCs and related pathogenic B cell populations in SLE are associated with distinct metabolic states (*9, 19, 30*). Here, transcriptomic analyses revealed broad suppression of OXPHOS-related genes after miR-1285-3p transfection, mirroring patterns observed in naïve B cells from patients with SLE. Functional assays demonstrated reduced mitochondrial mass and membrane potential, consistent with impaired mitochondrial activity. Furthermore, pharmacologic perturbation of mitochondrial metabolism functionally supported this link. Low-dose rotenone, which inhibits mitochondrial complex I, increased the frequency of ABCs, whereas activation of OXPHOS with AICAR suppressed their induction. In contrast, inhibition of ATP synthase with oligomycin did not phenocopy complex I inhibition (rotenone), suggesting that altered electron transport, rather than reduced ATP production per se, is more closely linked to this phenotype. A recent study showed that intact mitochondrial metabolism and OXPHOS activity are required to sustain TLR7-driven ABC differentiation by supporting mitochondrial quality control and cellular fitness (*31*). Taken together with that work, our findings suggest that ABC differentiation may depend on a tuned metabolic balance rather than a simple binary OXPHOS state, potentially reflecting context-dependent and stage-dependent requirements, as well as differential effects of partial versus more complete inhibition of mitochondrial respiration.

Among the transcriptional changes downstream of OXPHOS inhibition, IKZF2 emerged as a particularly compelling candidate regulator. IKZF2, also known as Helios, is a member of the Ikaros family and has been extensively studied in T cell biology, where it contributes to immune regulation and maintenance of regulatory T cell programs (*32, 33*). Although its function in B cells has remained poorly defined, human genetic studies have linked *IKZF2* to immune dysregulation and systemic autoimmunity, including SLE (*23, 24, 34–36*). In our study, *IKZF2* expression was significantly lower in selected B cell subsets from patients with SLE, including naïve B cells, DN B cells, and plasmablasts, whereas other B cell subsets showed no significant difference. This pattern indicates that *IKZF2* downregulation is not uniform across the B cell compartment but is enriched in subsets most relevant to disease-associated B cell differentiation. Consistent with these transcriptomic observations, IKZF2 (Helios) protein expression was decreased in circulating ABCs from patients with SLE. Notably, analysis of individuals carrying a germline IKZF2 loss-of-function mutation provided *in vivo* human genetic support for this model, as these individuals showed a skewing toward increased frequency of ABCs. Together, these data support the notion that reduced *IKZF2* expression is a disease-relevant feature of B cells in SLE.

Our functional data further support a regulatory role for IKZF2 in restraining ABC differentiation. Pharmacologic degradation of IKZF2 or siRNA-mediated knockdown was sufficient to promote ABC differentiation. Time-course analyses showed that suppression of IKZF2 preceded induction of *ITGAX* (CD11c) and was followed by delayed upregulation of *TBX21* (T-bet), consistent with a model in which loss of IKZF2 releases an early transcriptional brake before full acquisition of the ABC phenotype. In addition, degradation of IKZF2 attenuated the inhibitory effects of AICAR, supporting the idea that IKZF2 mediates the effects of OXPHOS-linked signaling on ABC differentiation. Thus, the OXPHOS–IKZF2 axis appears to act upstream of key ABC-associated transcriptional changes.

Our data also refine the stage at which IKZF2 is most relevant. Following IKZF2 degradation, *ITGAX* and *TBX21* were induced, whereas *ZEB2* remained unchanged despite detectable *IKZF2* occupancy at the *ZEB2* locus in public ChIP-seq data. This dissociation suggests that chromatin occupancy alone is not sufficient to predict transcriptional output and that additional context-dependent cofactors are likely required. More importantly, in the ImmuNexUT dataset, *IKZF2* expression in naïve B cells showed a moderate inverse correlation with *ITGAX* and *TBX21* within the same subset and a weaker but still significant inverse correlation with these genes in DN B cells, suggesting stage-specific regulation of ABC-associated gene programs. By contrast, *IKZF2* expression in DN B cells did not correlate with *ITGAX* or *TBX21*. This pattern argues that IKZF2 functions primarily at the naïve B cell stage, where it restrains acquisition of early ABC-associated gene programs, rather than acting as a dominant regulator after cells have already entered ABC-like state. Thus, IKZF2 appears to function less as a maintenance factor in differentiated ABCs and more as a gatekeeper of pathogenic B cell fate commitment.

These findings have potential translational implications. Small EV-associated miR-1285-3p may represent a potential biomarker reflecting upstream pathogenic activity linked to ABC expansion and clinical disease severity in SLE. Furthermore, the miR-1285-3p–OXPHOS–IKZF2 axis suggests several possible therapeutic entry points. Interventions that reduce pathogenic miRNA transfer, restore mitochondrial metabolic balance, or preserve IKZF2 function may limit ABC differentiation. Whether such approaches can selectively restrain pathogenic B cell programs without broadly compromising protective humoral immunity will require further study. Importantly, the proposed OXPHOS–IKZF2 axis was supported by multiple complementary lines of evidence, including in vitro perturbation assays, transcriptomic analyses of B cells, public epigenomic data, and independent human genetic evidence from IKZF2 loss-of-function mutation carriers. This convergence supports the biological and clinical relevance of this pathway in pathogenic B cell differentiation in SLE.

Several limitations should be acknowledged. First, our study focused on ABC differentiation, and the effects of miR-1285-3p and OXPHOS perturbation on other B cell subsets or immune cells remain to be defined. Second, the cellular sources of SLE-derived small EVs enriched for miR-1285-3p and the upstream mechanisms that regulate this enrichment are still unknown. Third, although our data support a mechanistic pathway linking miR-1285-3p, OXPHOS suppression, and reduced IKZF2 activity, *in vivo* validation will be necessary to establish the contribution of this axis to lupus pathogenesis. The analysis of individuals carrying a germline IKZF2 loss-of-function mutation was limited by the small sample size and the observed ABC expansion may partly reflect indirect effects mediated by alterations in other immune cells beyond B cells (*24*). Nevertheless, these in vivo findings provided an important proof-of-concept supporting the disease relevance of IKZF2 in human autoimmunity. Finally, although our study centered on SLE, ABC-like populations expand across multiple autoimmune diseases, and it remains unclear whether the miR-1285-3p-OXPHOS-IKZF2 axis is specific to SLE or reflects a broader mechanism of pathogenic B cell differentiation across autoimmunity.

In summary, our study identifies circulating small EV–associated miR-1285-3p as a potential upstream modulator of pathogenic B cell differentiation in SLE and establishes an OXPHOS–IKZF2 axis that restrains early ABC-associated gene programs. These findings provide a mechanistic framework linking extracellular miRNA signaling to the metabolic and transcriptional control of B cell fate in human autoimmunity.

## Materials and Methods

### Human subjects

Human samples and corresponding clinical information were collected from the University of Tsukuba Hospital in Japan and from a previously reported cohort (*24*). Patients were diagnosed with SLE according to the 2019 European League Against Rheumatism/American College of Rheumatology classification criteria for systemic lupus erythematosus (*37*). Patients with SLE and healthy controls (HCs) provided informed consent under ethical and safety protocols approved by the ethics committee of the University of Tsukuba Hospital (No. H24-164) and in accordance with the original study (*24*). Written informed consent was obtained from all patients before participation in the study. Participant information is provided in Table S1.

### Naïve B cell preparation

Peripheral blood mononuclear cells (PBMCs) were isolated from donors using Ficoll-Paque PLUS (GE Healthcare). CD19^+^IgD^+^CD27^-^ B cells were isolated from PBMCs using the EasySep^TM^ Human Naïve B Cell Isolation Kit (purity >90%) (STEMCELL Technologies), according to the manufacturer’s instructions.

### Flow cytometry

For surface staining, single-cell suspensions were prepared from PBMCs or B cells and stained with fixable viability dye for 30 min at 4°C, followed by staining with surface antibodies for 30 min at 4°C. Antibodies used for flow cytometry are listed in Table S2. For intracellular staining of transcription factors, including IKZF2 (Helios) or T-bet, cells were fixed and permeabilized using the eBioscience™ Foxp3 / Transcription Factor Staining Buffer Set (Thermo Fisher Scientific), according to the manufacturer’s instructions. After fixation and permeabilization, cells were stained with intracellular antibodies for 30 min at 4°C, washed, and resuspended in staining buffer prior to acquisition. For assessment of mitochondrial mass, membrane potential, or mitochondrial reactive oxygen species (mtROS), cells were resuspended in prewarmed culture medium containing MitoSpy dyes or MitoSOX Red after viability staining and incubated for 30 min at 37°C.

Cells were analyzed using the BD LSRFortessa™ X-20 Flow Cytometer (BD Biosciences) and FlowJo software (Tree Star).

### Isolation of small EVs

Small EVs were isolated from plasma by ultracentrifugation according to previously described protocols (*38*). In brief, plasma from patients with SLE or HCs was centrifuged sequentially at 300g for 10 min, 2,000g for 10 min and 10,000g for 30 min at 4°C. The supernatants were filtered through a 0.2 μm pore-size filter (Sartorius) to remove large EVs such as apoptotic bodies and microvesicles, then ultracentrifuged at 110,000g for 2 hours at 4°C three times. Pellets were resuspended in phosphate-buffered saline (PBS) and stored at −80°C. The size distribution of collected small EVs was measured using the NanoSight LM10 (Malvern Instruments). In addition, transmission electron microscopy (TEM) was performed to confirm the morphology of the isolated small EVs by the Hanaichi Ultrastructure Research Institute (Okazaki).

### Western blotting

Proteins were extracted from small EVs and cells using 5× sample buffer containing 2-Mercaptoethanol (0.5 M Tris-HCl [pH 6.8], 100% Glycerol, 10% SDS, and Bromophenol Blue) supplemented with Protease/ Phosphatase Inhibitor Cocktail (Cell Signaling Technology). Extracted proteins were loaded onto SuperSep (TM) Ace Gel, 5-20% (FUJIFILM Wako), incubated for 5 min at 95°C, and separated by electrophoresis (25 mA for 65 min). Proteins were transferred to polyvinylidene fluoride membranes (Bio-Rad), according to the manufacturer’s instructions. Membranes were blocked in 1× TBS (pH 7.4) (NIPPON GENE) containing 0.1% Tween 20 (MP Biomedicals) and 5% skim milk (FUJIFILM Wako) for 1 hour at room temperature, followed by incubation overnight at 4°C with the following primary antibodies. Primary and secondary antibodies used for immunoblotting are listed in Table S2. Membranes were then incubated with HRP-conjugated secondary antibodies for 1 hour at room temperature. Digital images were obtained using the FUSION FX7 (Vilber Bio Imaging) with Western Chemiluminescent HRP substrate (Cytiva).

### Isolation of miRNA and mRNA

For circulating miRNA analysis, miRNAs were isolated from circulating small EVs using the microRNA Extractor® Kit for Purified EV (FUJIFILM Wako), according to the manufacturer’s instructions. As an exogenous control, 0.1 pM synthetic cel-miR-39-3p RNA oligonucleotide (UCACCGGGUGUAAAUCAGCUUG) (Thermo Fisher Scientific) was added to each sample after RNA extraction. For cultured cell experiments, miRNAs were isolated from harvested cells using Isogen II (NIPPON GENE) according to the manufacturer’s instructions.

Total RNA, including messenger RNA (mRNA), was isolated from cultured cells using the microRNeasy Micro Kit Plus (QIAGEN), according to the manufacturer’s instructions.

### Quantitative real-time PCR (qPCR) for miRNA and mRNA

For circulating miRNAs derived from small EVs and miRNAs from cultured cells, complementary DNA (cDNA) was synthesized using the TaqMan™ MicroRNA Reverse Transcription Kit (Applied Biosystems). Quantitative real-time PCR was performed using TaqMan™ Universal PCR Master Mix (Thermo Fisher Scientific) and the StepOnePlus™ Real-Time PCR System (Thermo Fisher Scientific). Expression levels of circulating miRNAs were calculated using the 2^−ΔCt^ method and normalized to spiked-in cel-miR-39-3p. To assess miRNA transfection efficiency in cultured cells, qPCR data were also analyzed using the 2^−ΔΔCt^ method, with miR-24-3p as an endogenous normalization control. TaqMan™ MicroRNA Assays used in this study are listed in Table S2.

For mRNA expression analysis, cDNA was synthesized from total RNA using PrimeScript™ RT Master Mix (Perfect Real Time) (Takara Bio), according to the manufacturer’s instructions. qPCR was performed using TaqMan™ Universal PCR Master Mix and the StepOnePlus™ Real-Time PCR System. Gene expression levels were calculated using the 2^−ΔΔCt^ method and normalized to beta-2-microglobulin (B2M). TaqMan™ Gene Expression Assays used in this study are listed in Table S2.

### Cell culture

Naïve B cells from HCs were seeded at 1.0 × 10^5^ cells per well in a 96-well plate with RPMI-1640 medium (FUJIFILM Wako) supplemented with 10% fetal bovine serum and 1% penicillin/streptomycin (Gibco). Cells were incubated at 37°C in a humidified atmosphere containing 5% CO₂. To induce ABC differentiation, cells were stimulated for 72 hours with a cytokine cocktail containing 20 ng/mL IFN-γ, 1 μg/mL R848 (TLR7 agonist), 55 ng/mL IL-2, 10 ng/mL IL-21, 10 ng/mL BAFF, and 10 μg/mL anti-IgG (*8*). For the subsequent 72 hours, cells were restimulated with 20 ng/mL IFN-γ, 1 μg/mL R848, and 10 ng/mL IL-21.

0.1 μM rotenone (a mitochondrial complex I inhibitor and OXPHOS inhibitor), 3 μM oligomycin (an ATP synthase inhibitor), 100 μM AICAR (an AMP-activated protein kinase agonist and OXPHOS activator), or 5 mM N-acetylcysteine was added 30 min prior to cytokine stimulation, followed by incubation under the same culture conditions. NVP-DKY709 (4 μM), a selective IKZF2 degrader, was added simultaneously with the cytokine cocktail. In parallel, the human B cell line Raji (JCRB Cell Bank, JCRB9012) was cultured under the same conditions and seeded at 1.0 × 10⁵ cells per well in a 48-well plate. Cells were cultured for 72 hours in the presence or absence of ABC-inducing conditions, with or without rotenone (0.1 μM), which was added 30 min before stimulation. Unlike primary B cells, Raji cells were cultured without restimulation.

Cytokines, pharmacologic reagents, and antibodies used in this study are listed in Table S2.

### Microarray analysis of miRNAs

RNA was extracted from small EVs using 3D-Gene RNA extraction reagent from liquid sample (Toray Industries), according to the manufacturer’s instructions. Extracted total RNA was assessed using the Bioanalyzer (Agilent Technologies) and labeled with 3D-Gene miRNA labeling kit (Toray Industries). Half the volume of labeled RNA was hybridized onto the 3D-Gene (Human) miRNA Oligo chip (Toray Industries). The probe annotation and oligonucleotide sequences were based on the miRBase miRNA database (http://microrna.sanger.ac.uk/sequences/). After stringent washes, fluorescent signals were scanned with the 3D-Gene Scanner (Toray Industries) and analyzed using 3D-Gene Extraction software (Toray Industries).

The raw data for each spot were normalized by substitution with the mean intensity of the background signal determined from the signal intensities of all blank spots within the 95% confidence intervals. Measurements of spots with signal intensities greater than 2 standard deviations (SD) above the background signal intensity were considered valid. The relative expression level of a given miRNA was calculated by comparing the signal intensities of valid spots across the microarray experiments. The normalized data were globally normalized for each array such that the median signal intensity was adjusted to 25.

### Transfection of miRNA mimics or siRNA

Human naïve B cells, prepared as described above, were transfected with miRNA mimics, siRNA, or negative control (NC) using the 4D-Nucleofector^TM^ X Unit and P3 kit (Lonza). The final concentration was set at 6 pM. Cells were rested overnight after transfection and then seeded at 2.0 × 10^5^ cells per well, stimulated with 20 ng/mL IFN-γ, 1 μg/mL R848, 55 ng/mL IL-2, 10 ng/mL IL-21, 10 ng/mL BAFF, and 10 μg/mL anti-IgG, and cultured for 72 hours. For the subsequent 72 hours, cells were restimulated with 20 ng/mL IFN-γ, 1 μg/mL R848, and 10 ng/mL IL-21 prior to flow cytometric analysis. The sequences of miRNA mimics used in this study were as follows: miR-1285-3p (UCUGGGCAACAAAGUGAGACCU) and miR-619-5p (GCUGGGAUUACAGGCAUGAGCC). All miRNA mimics, siRNAs, and control reagents are listed in Table S2.

### Bulk RNA-seq

After total RNA was extracted using microRNeasy Micro Kit Plus (QIAGEN), mRNA was isolated using oligo magnetic beads and fragmented for cDNA synthesis. RNA integrity was assessed using the Bioanalyzer 2100 system (Agilent Technologies). Libraries were generated using the NEBNext Ultra^TM^ RNA Library Prep Kit (New England Biolabs) for the Illumina system according to the manufacturer’s instructions. Sequencing was conducted using the Illumina HiSeq X Ten platform.

### RNA-seq data analysis

Low quality ends (Phred score <30) and short reads (minimum length = 30) were trimmed using PRINSEQ (v0.20.4) (*39*). Trimmed reads were aligned to the hg38 reference genome using STAR (v2.7.8a) (*40*), and RSEM (v1.3.3) (*41*) was used to quantify reads mapping to genes from Ensembl release 114. Pairwise differential expression analysis was performed using the R package DESeq2 (*42*). RNA-seq data will be deposited in the NCBI Gene Expression Omnibus (GEO) database.

Public transcriptomic data from the ImmuNexUT cohort were obtained from the National Bioscience Database Center (NBDC; JGAS000486) (*20*). Raw read counts were normalized using the trimmed mean of M-values (TMM) method implemented in the R package edgeR (v4.8.2), followed by transformation to counts per million (CPM). Expression values were log-transformed as log(CPM + 1). Batch effects were corrected using the ComBat function from the R package sva (v3.58.0).

Pathway enrichment analysis was performed using Enrichr with the WikiPathways 2024 Human database. Downregulated genes were defined as those with *P* < 0.1 and log_2_ fold change < −0.5 in bulk RNA-seq data from miR-1285-3p-transfected B cells.

### IKZF2 ChIP-seq analysis in human GM12878 cells

Publicly available ChIP-seq data targeting *IKZF2* in human GM12878 cells were obtained from the ENCODE Project (GSE105865), and input ChIP-seq data from GM12878 cells (GSE92188) were analyzed as controls. Paired-end reads generated on the Illumina HiSeq 4000 platform were subjected to adapter trimming using Trim Galore (version 0.6.10), followed by alignment to the human reference genome (hg38) using Bowtie2 (version 4.1.2). Aligned reads with a mapping quality score (MAPQ) ≥30 were retained, and PCR duplicates and blacklisted genomic regions defined by the ENCODE consortium were removed. The resulting reads were then sorted and indexed using SAMtools (version 1.8). ChIP-seq peaks were identified using MACS2 (version 2.2.7.1) with the callpeak command and the following parameters: --format BAMPE --keep-dup all --gsize hs --qvalue 0.05. After filtering, read coverage tracks were generated by converting aligned reads into bigWig format. The resulting signal tracks were visualized together with ChIP-seq peaks using the Integrative Genomics Viewer (IGV) (version 2.16.0). Peaks were assigned to the nearest genes based on distance to transcription start sites (TSSs) using the R package ChIPseeker (version 1.30.3) and were classified according to genomic features, including promoter, exon, intron, and intergenic regions.

### Statistical analysis

Data were analyzed using FlowJo (BD Biosciences, USA), GraphPad Prism (GraphPad Software, CA, USA), and R version 4.4.1 (R Core Team). Probability values below 0.01, 0.05, or 0.1 were considered significant, as specified for individual analyses.

## Acknowledgments

We would like to thank the staff at Department of Rheumatology, Institute of Medicine, University of Tsukuba for their dedicated efforts in supporting this research. We also thank Cosmin Mihail Florescu (Medical English Communications Center, University of Tsukuba) for his editorial assistance.

## Funding

This work was also supported by JST SPRING, Grant Number JPMJSP2124.

## Author contributions

S.A. performed experiments, analyzed data, and wrote the original draft. S.A. and H.A. designed the study. H.I. and K.I. performed ChIP-seq analyses. H.A., T.A., and M.O. performed bulk RNA-seq analyses. N.U., H.M., T.N., A.O., Y.K., H.T., and I.M. helped with sample collection and experiments. K.F. and I.M. supervised the experiments. All authors reviewed the manuscript.

## Competing interests

The authors declare that they have no competing interests.

## Data and materials availability

Data-sharing requests, which should be directed to the corresponding author, will be reviewed upon request.

## Supplementary Materials

**Supplementary Fig. 1. Flow cytometry gating strategy for defining ABCs.**

Following exclusion of doublets and dead cells, IgD⁻CD27⁻ double-negative (DN) B cells were gated. From this DN population, CD21⁻ cells were selected, and ABCs were defined as the CD11c⁺T-bet⁺ subset within the DN CD21⁻ B-cell fraction.

**Supplementary Fig. 2. Transfection efficiency of miR-1285-3p and miR-619-5p mimics in naïve B cells.**

Relative expression of miR-1285-3p and miR-619-5p in B cells 48 hours after transfection with each miRNA mimic or negative control (NC) (n = 4 per group). Data are presented as mean ± SEM. \**P* < 0.05; Mann–Whitney U test.

**Supplementary Fig. 3. OXPHOS-related pathways are enriched among downregulated genes in naïve B cells from patients with SLE.**

Pathway enrichment analysis of genes downregulated in naïve B cells from patients with SLE using the publicly available dataset GSE92387. Pathways are ranked by −log₁₀(*P* value).

**Supplementary Fig. 4. Inhibition of ATP synthase with oligomycin does not alter ABC differentiation.**

Naïve B cells were cultured under ABC-inducing conditions and treated with the ATP synthase inhibitor oligomycin or DMSO. Quantification of ABC frequencies showed no significant difference between groups (n = 7 per group). ns, not significant; Wilcoxon signed-rank test.

**Supplementary Fig. 5. Rotenone enhances CD11c expression in Raji cells under basal and ABC-inducing conditions.**

(**A**) Representative histograms and quantification of CD11c expression (mean fluorescence intensity [MFI]) in Raji cells treated with rotenone or DMSO for 72 hours. (**B**) Representative histograms and quantification of CD11c expression (MFI) in Raji cells cultured under control conditions (DMSO), ABC-inducing conditions, or ABC-inducing conditions with rotenone for 72 hours. MFI values were calculated from viable Raji cells after exclusion of dead cells. Data are presented as mean ± SEM. \*\**P* < 0.01. Mann-Whitney U test was used for (**A**); Kruskal–Wallis test was used for (**B**).

**Supplementary Fig. 6. Acute mitochondrial ROS generation is not detected following complex I inhibition, whereas ROS scavenging partially attenuates rotenone-induced ABC differentiation.**

(**A**) Mitochondrial reactive oxygen species (mtROS) levels assessed by MitoSOX staining in naïve B cells cultured under ABC-inducing conditions and treated with DMSO, rotenone, or oligomycin for 1 hour (n = 4 per group), 2 hours (n = 6 per group), and 4 hours (n = 6 per group). (**B**) Frequencies of ABCs in B cells treated with DMSO, rotenone, or rotenone in combination with N-acetylcysteine (NAC) (n = 7 per group). Data are presented as mean ± SEM. \**P* < 0.05, \*\**P* < 0.01; ns, not significant. Kruskal–Wallis test with multiple comparisons was used for (**A**); Wilcoxon signed-rank test was used for (**B**).

**Supplementary Fig. 7. Validation of *IKZF2* downregulation following OXPHOS inhibition.**

qPCR analysis of *IKZF2* expression in B cells treated with rotenone or DMSO for 24 hours under ABC-inducing conditions (n = 6 per group). Data are presented as mean ± SEM. \*\**P* < 0.01; Mann–Whitney U test.

**Supplementary Fig. 8. Time-course analysis of *TBX21* expression following OXPHOS inhibition during ABC differentiation.**

qPCR analysis of *TBX21* expression in B cells treated with rotenone or DMSO under ABC-inducing conditions at the indicated time points (n = 6 per group). Data are presented as mean ± SEM. \**P* < 0.05; Mann-Whitney U test.

**Supplementary Fig. 9. Time-course analysis of *ZEB2* expression following OXPHOS inhibition during ABC differentiation.**

qPCR analysis of *ZEB2* expression in B cells treated with rotenone or DMSO under ABC-inducing conditions at the indicated time points (n = 6 per group). Data are presented as mean ± SEM. ns, not significant; Mann-Whitney U test.

**Supplementary Fig. 10. IKZF2 knockdown enhances ABC differentiation.**

(**A**) Representative flow cytometry plots showing frequencies of ABCs (CD11c⁺T-bet⁺) in naïve B cells transfected with IKZF2-targeting siRNA (siIKZF2) or control siRNA (siCtrl) under ABC-inducing conditions. (**B**) Quantification of ABC frequencies in B cells treated with siIKZF2 or siCtrl (n = 7 per group). Data are presented as mean ± SEM. \**P* < 0.05; Wilcoxon signed-rank test.

**Supplementary Fig. 11. *ZEB2* expression is not altered by IKZF2 degradation despite IKZF2 binding at the *ZEB2* locus.**

(**A**) Time-course qPCR analysis of *ZEB2* expression in B cells treated with NVP-DKY709 or DMSO under ABC-inducing conditions (n = 6 per group). (**B**) Representative genome browser tracks showing IKZF2 ChIP-seq signal, input control, and called peaks across the *ZEB2* locus in the human B cell line GM12878.

**Supplementary Fig. 12. Correlation between *IKZF2* expression and ABC-associated genes in DN B cells.**

Correlation between *IKZF2* expression and *ITGAX* or *TBX21* expression in DN B cells from patients with SLE. Each dot represents an individual sample (n = 134). Pearson’s correlation.

**Supplementary Fig. 13. Distribution of B cell subsets in individuals carrying an IKZF2 mutation.**

Frequencies of naïve, unswitched memory (USM), switched memory (SM), and double-negative (DN) B cells among CD19⁺ B cells in individuals carrying the IKZF2 mutation (P1 and P2) and HCs. Each dot represents an individual sample. Horizontal bars indicate the median. IKZF2-mutant, individuals carrying the IKZF2 mutation; HCs, healthy controls.

**Table S1.** Clinical characteristics of study participants

**Table S2.** Antibodies, reagents, and assay details used in this study

## Notes

### Competing Interest Statement

The authors have declared no competing interest.

