## Supplemental Figures for "Circulating miR-1285-3p promotes age-associated B cell differentiation through the OXPHOS-IKZF2 axis in SLE"

### Supplementary Figure 1

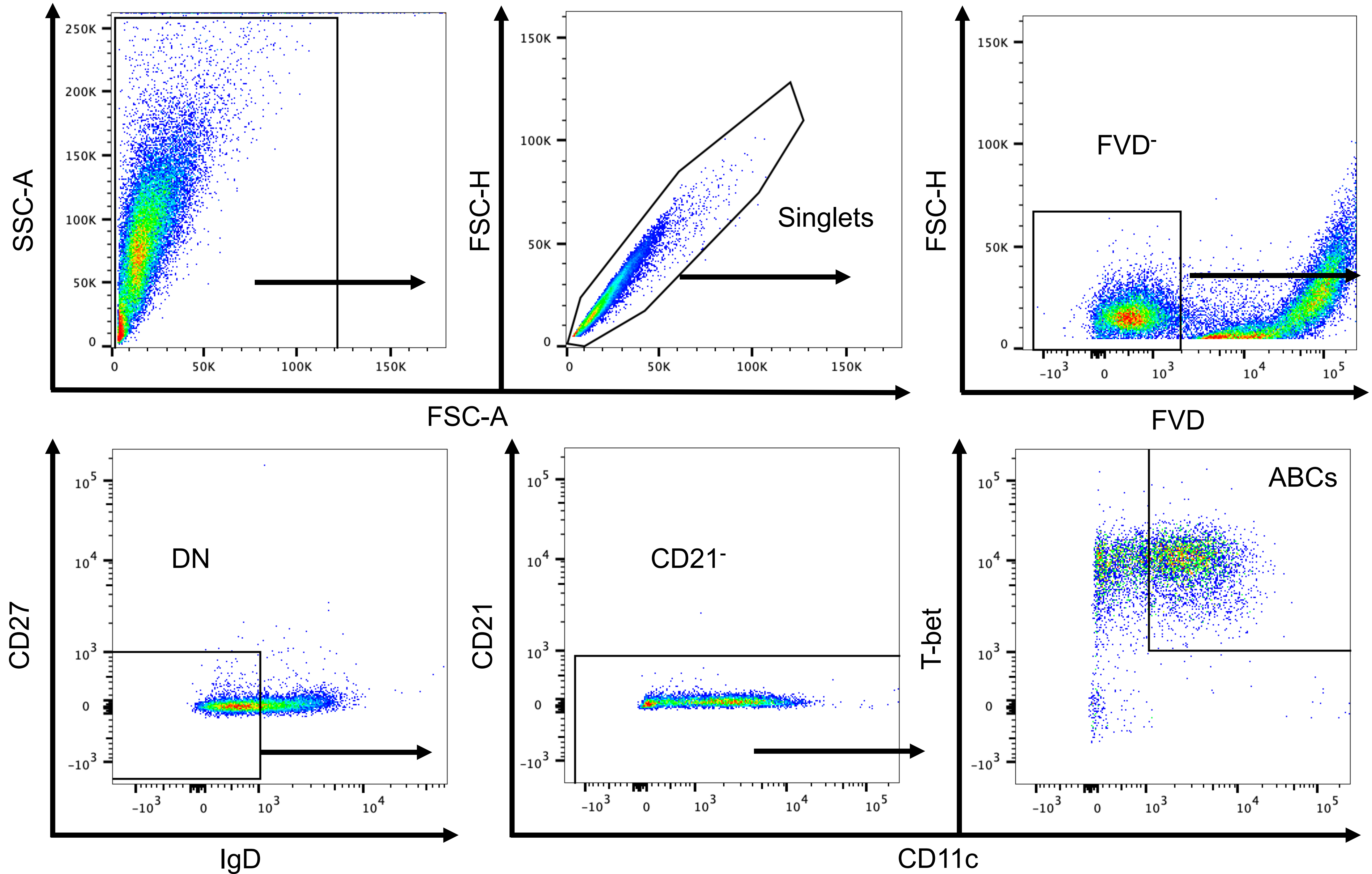

Supplementary Figure 2

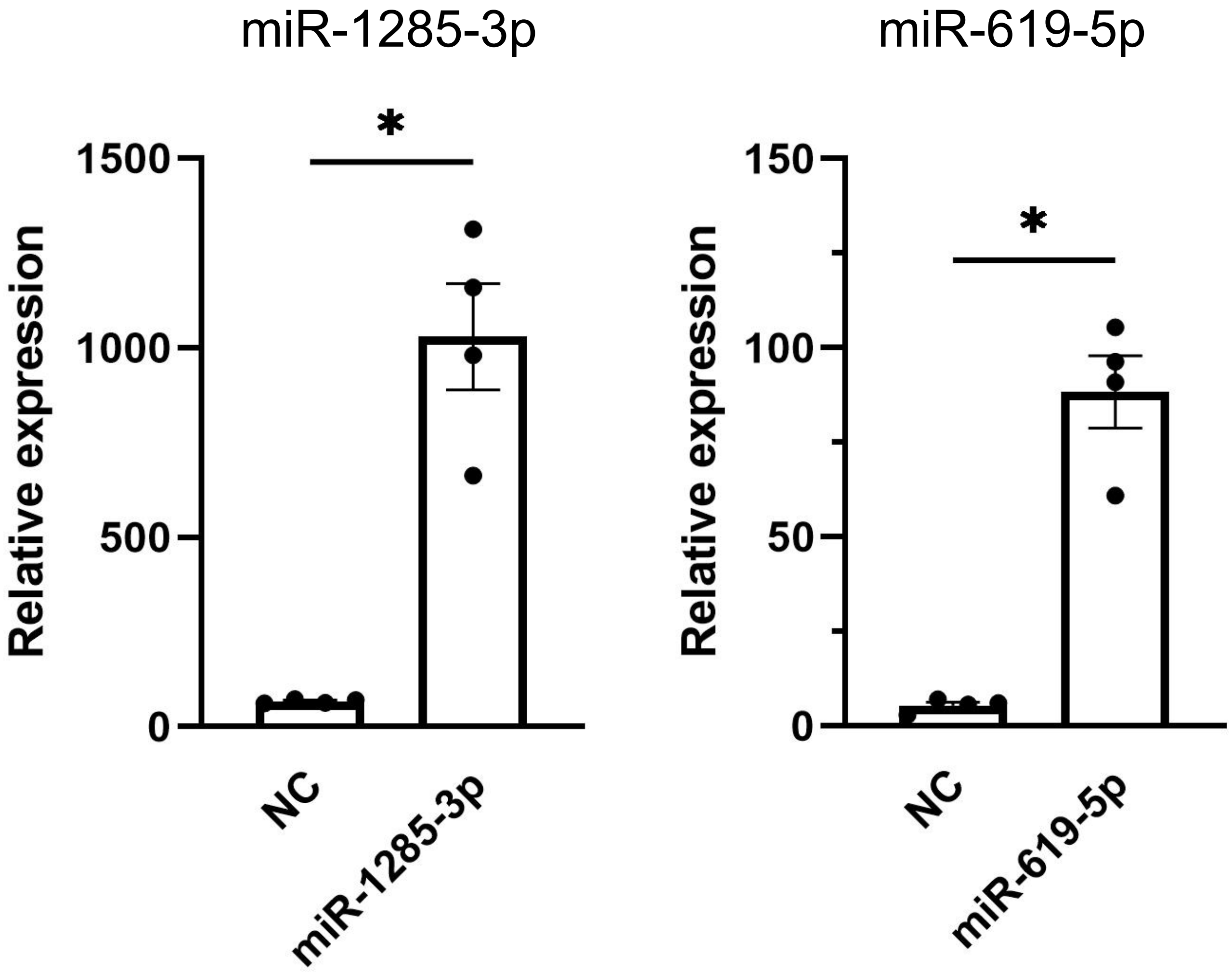

### Supplementary Figure 3

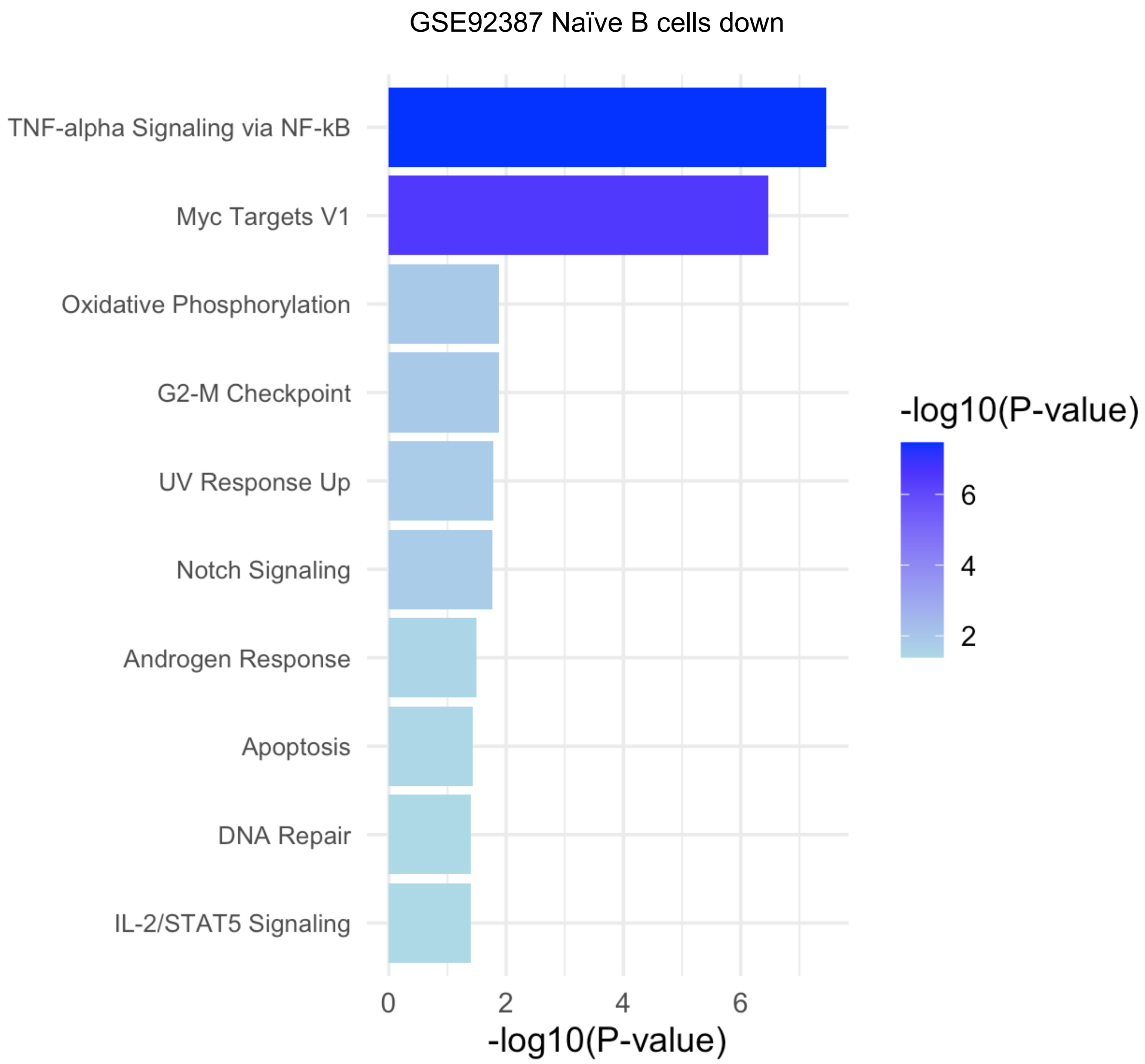

Supplementary Figure 4

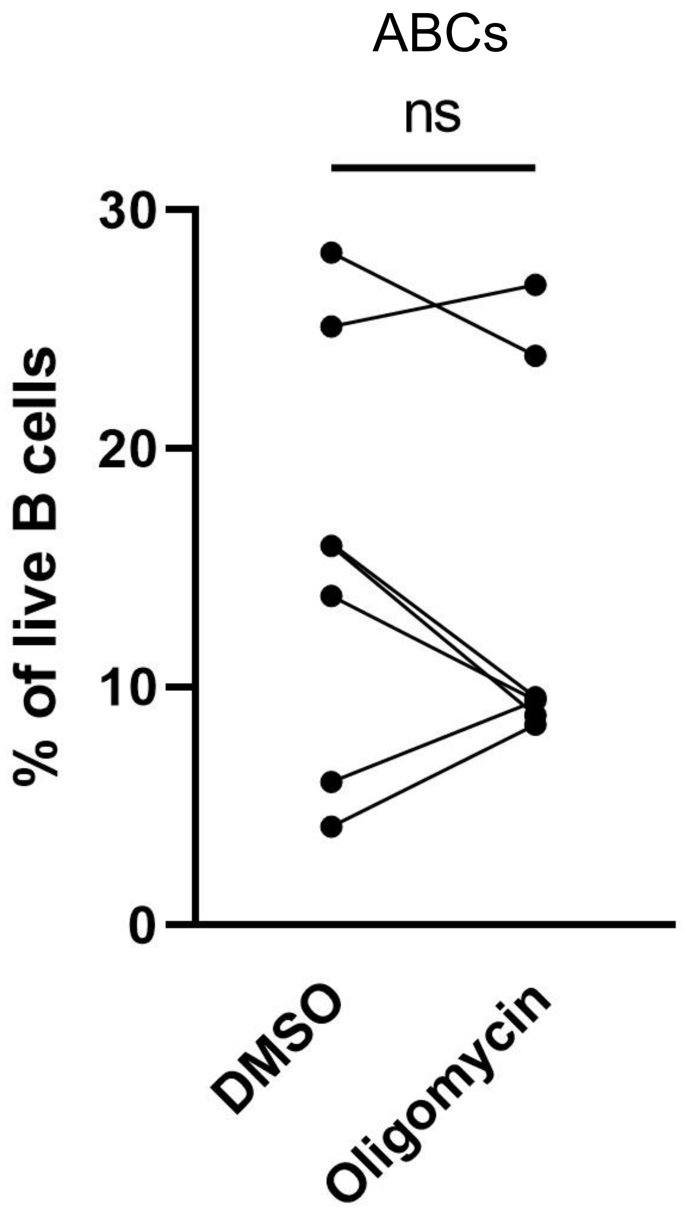

Supplementary Figure 5

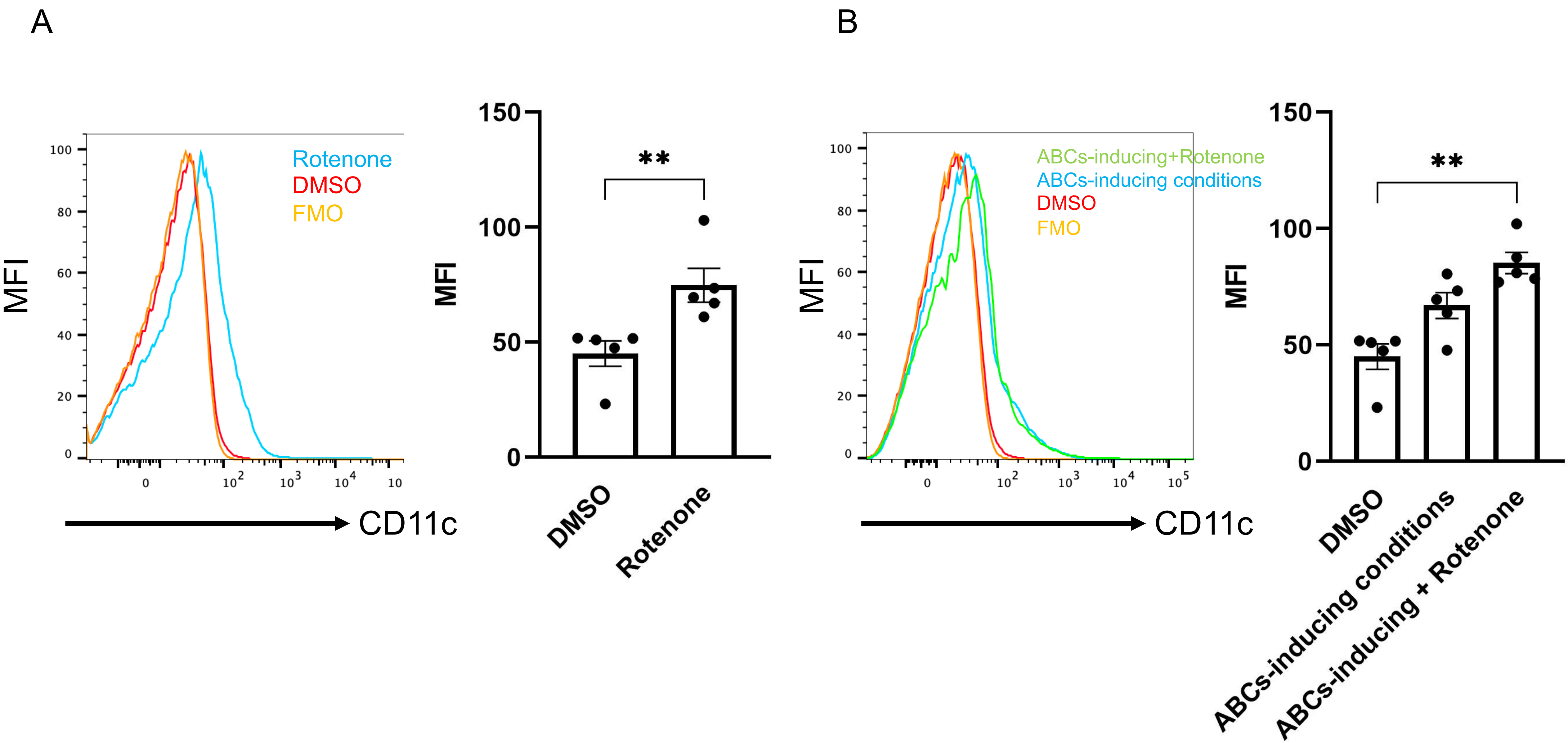

Supplementary Figure 6

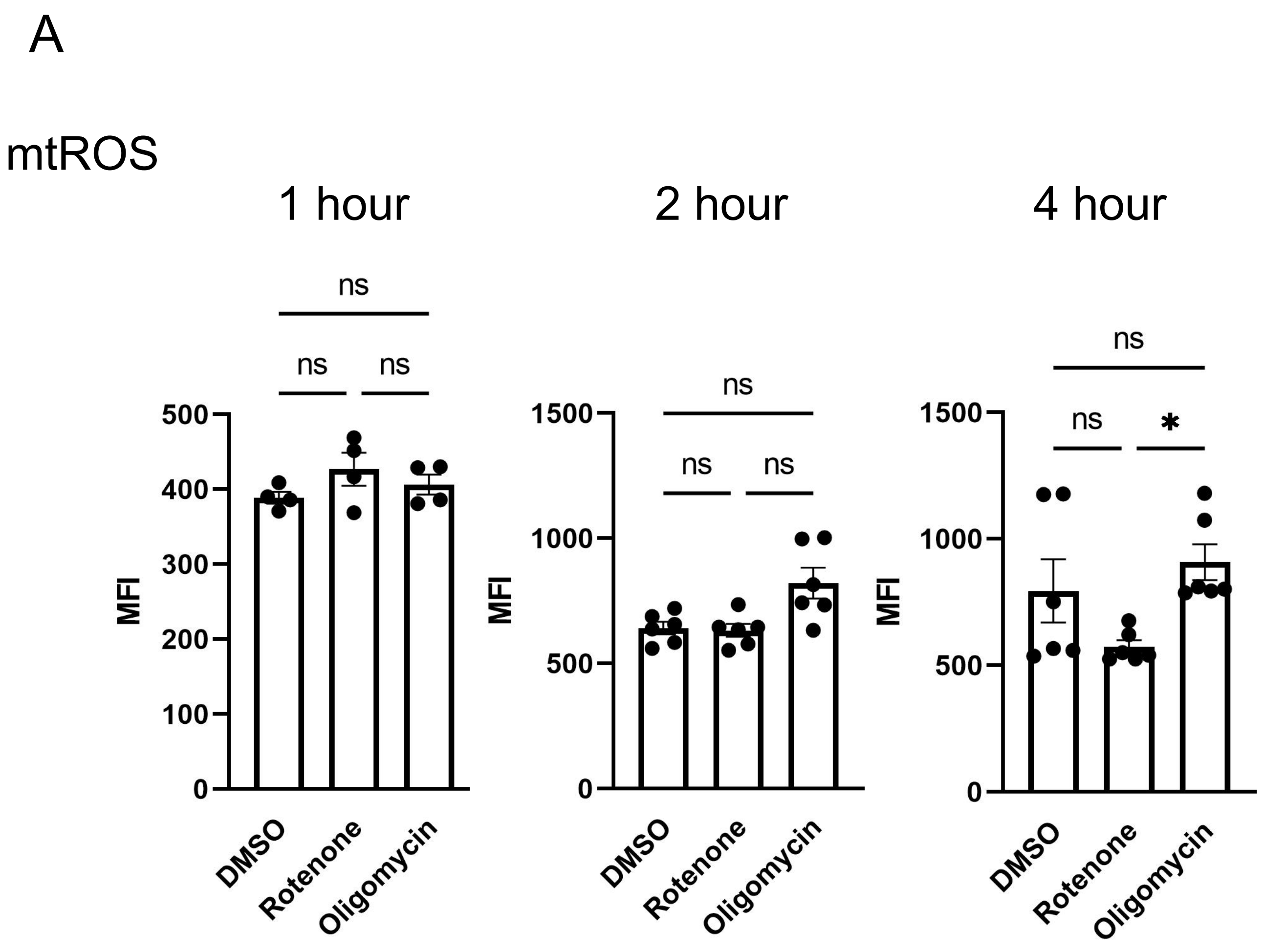


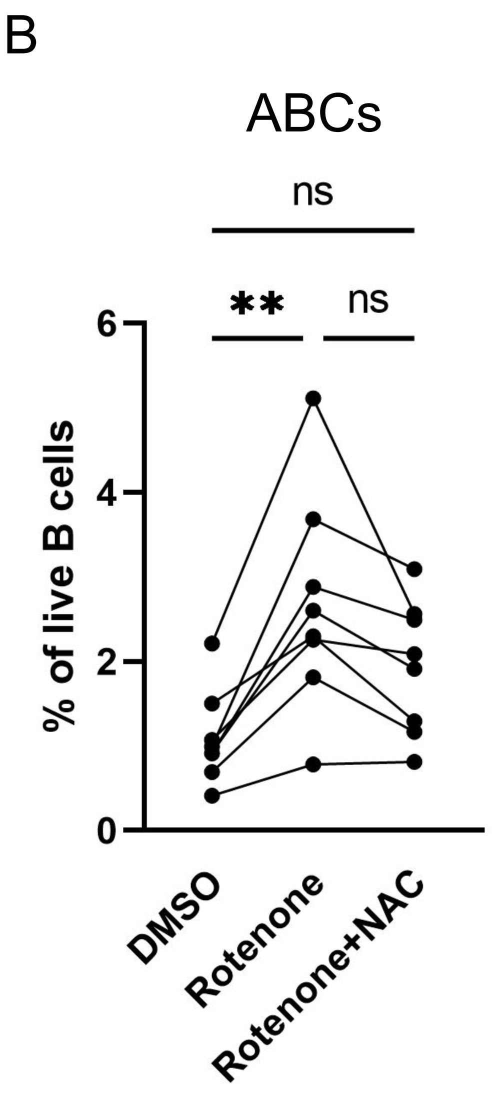

Supplementary Figure 7

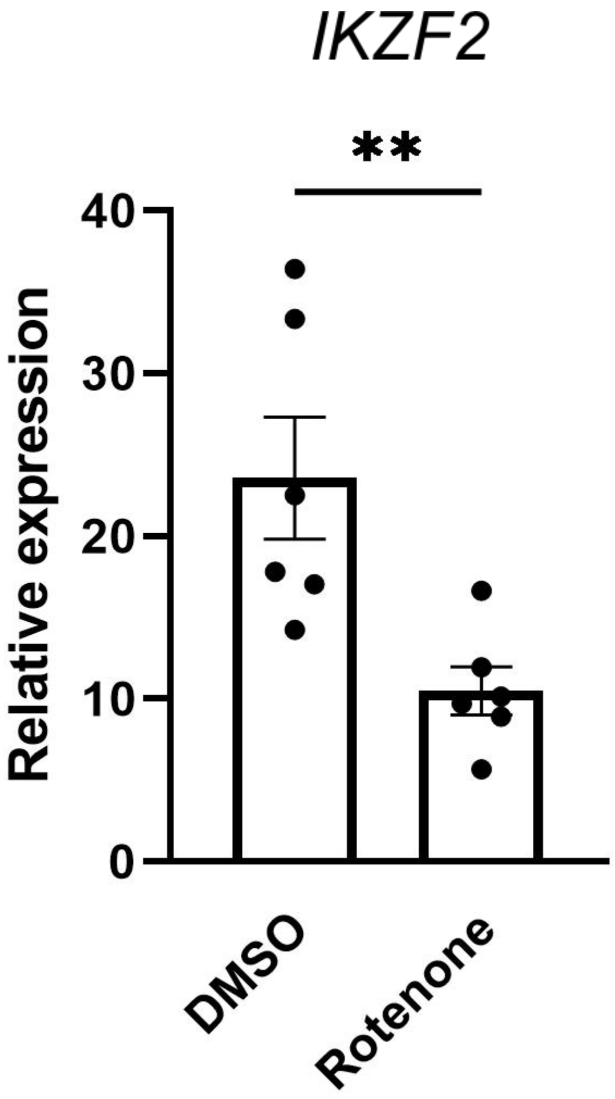

Supplementary Figure 8

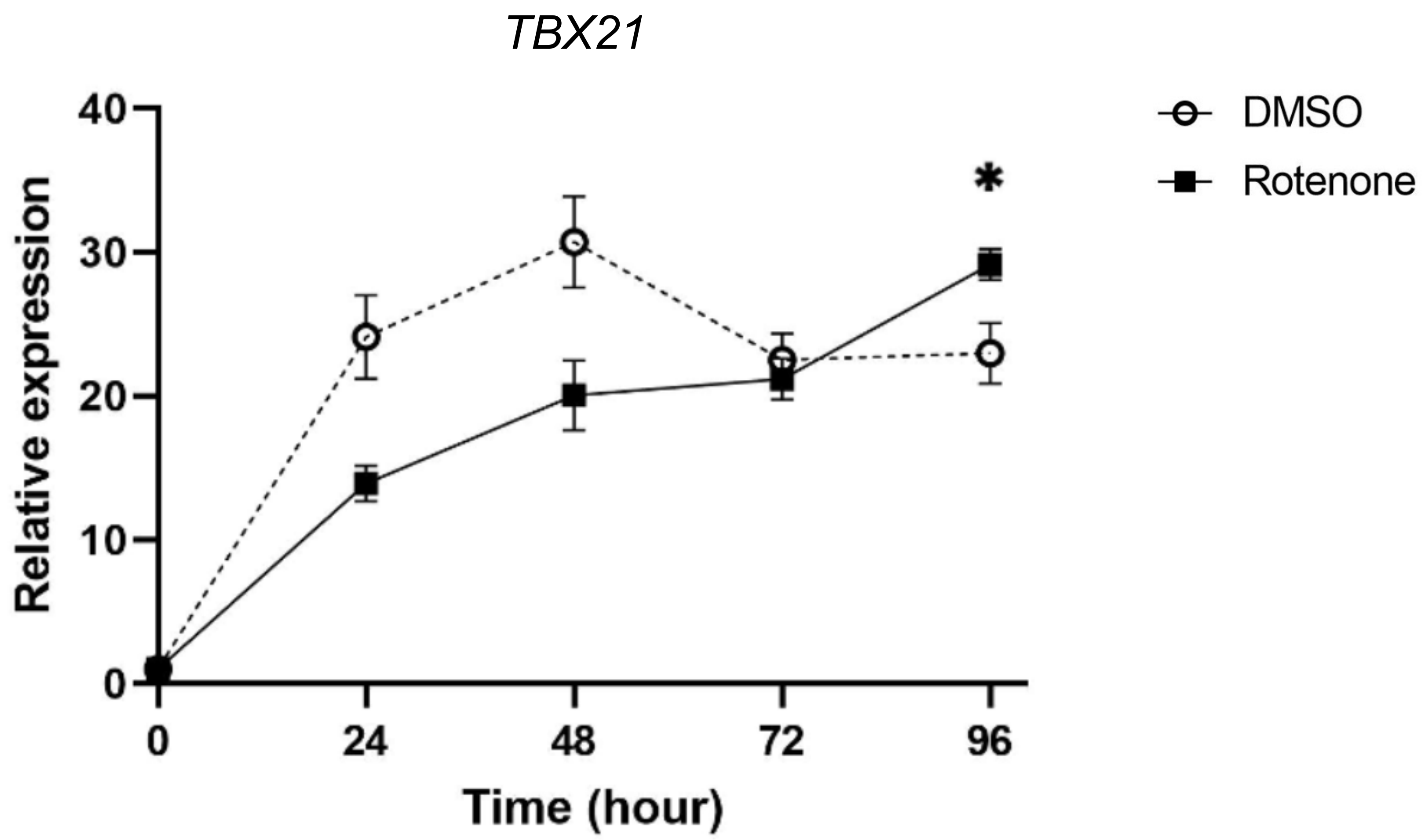

Supplementary Figure 9

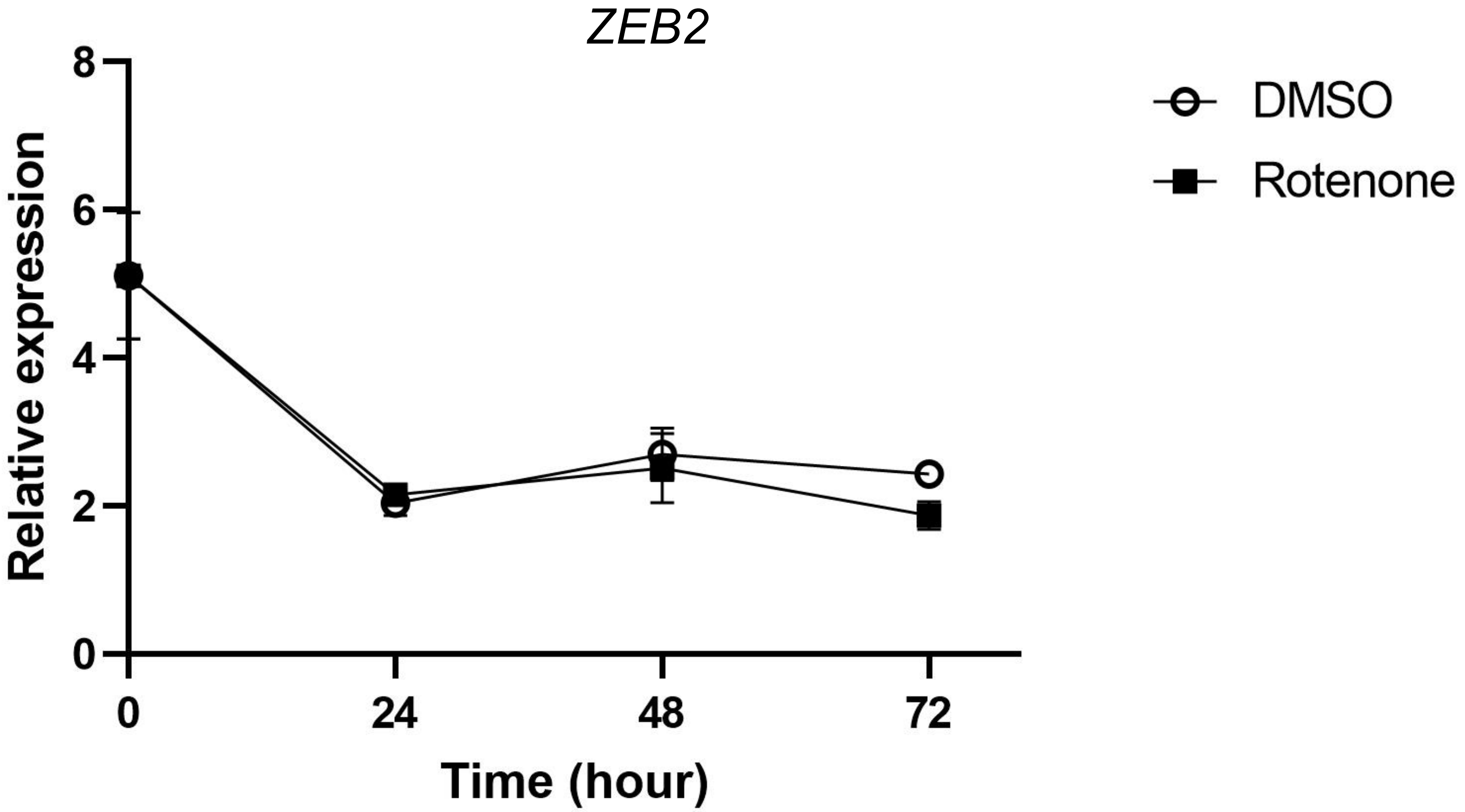

Supplementary Figure 10

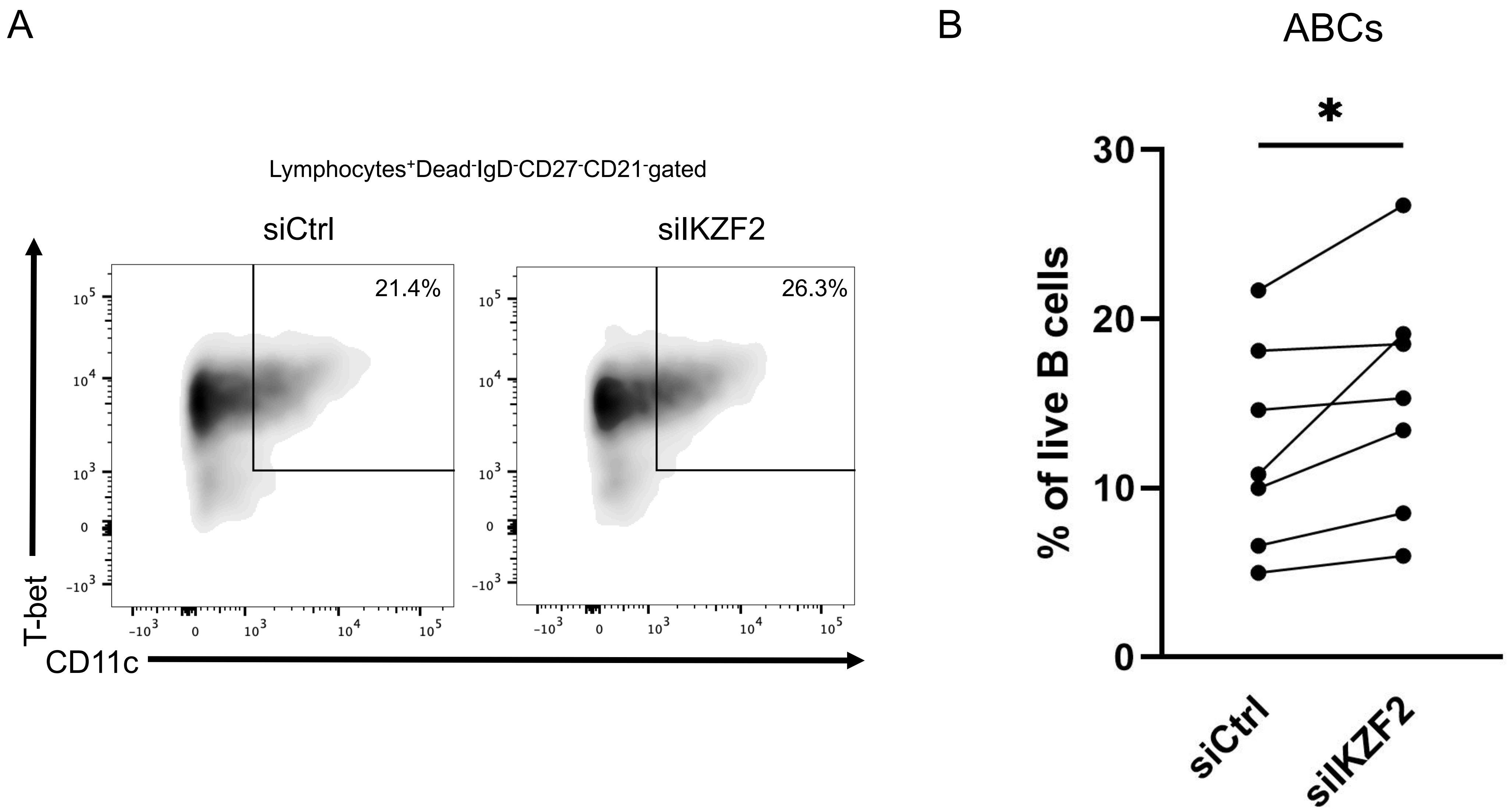

Supplementary Figure 11

A

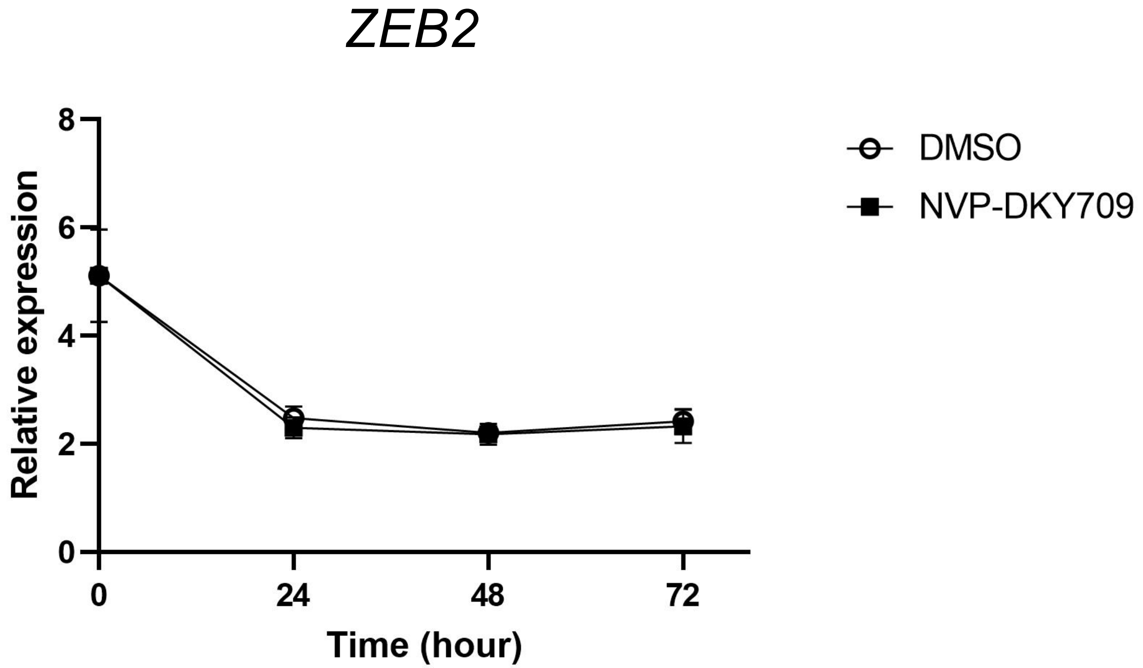


B ZEB2

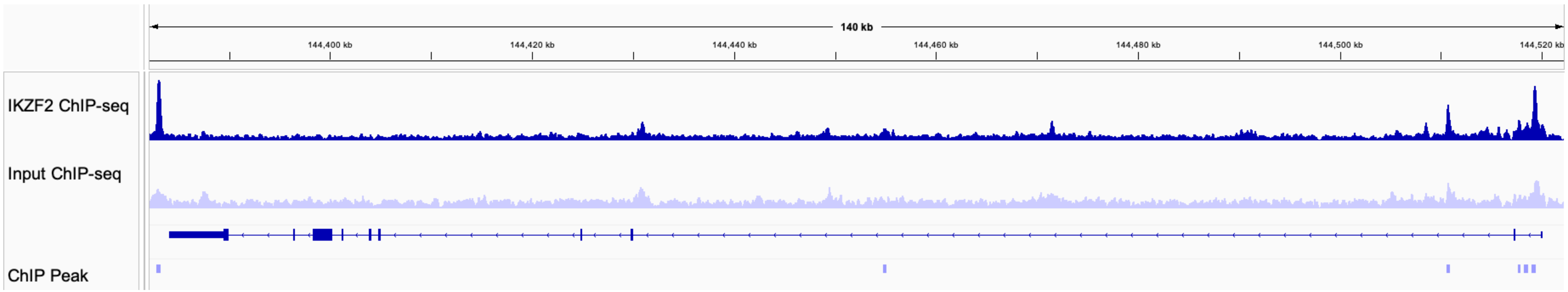

Supplementary Figure 12

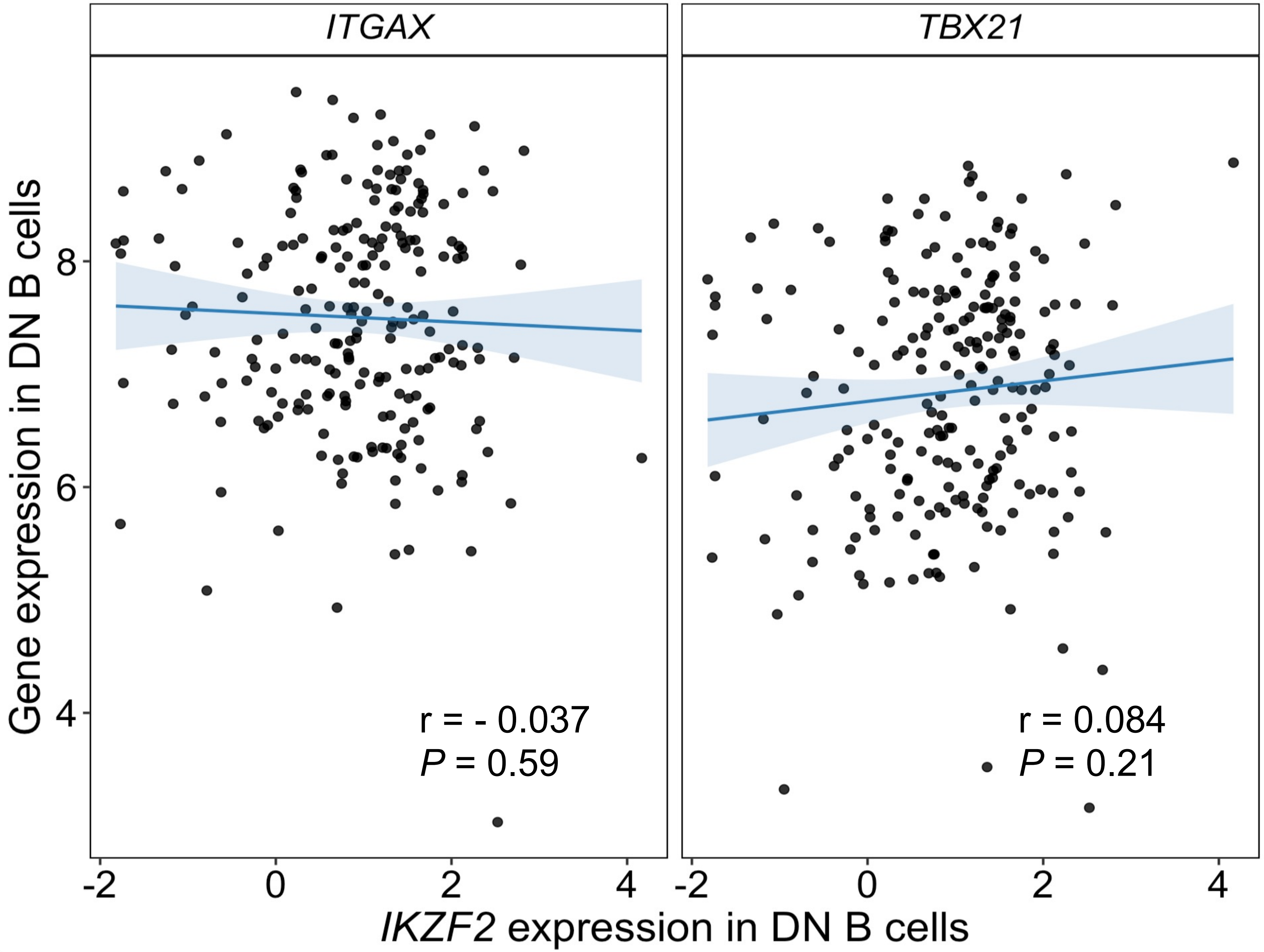

Supplementary Figure 13

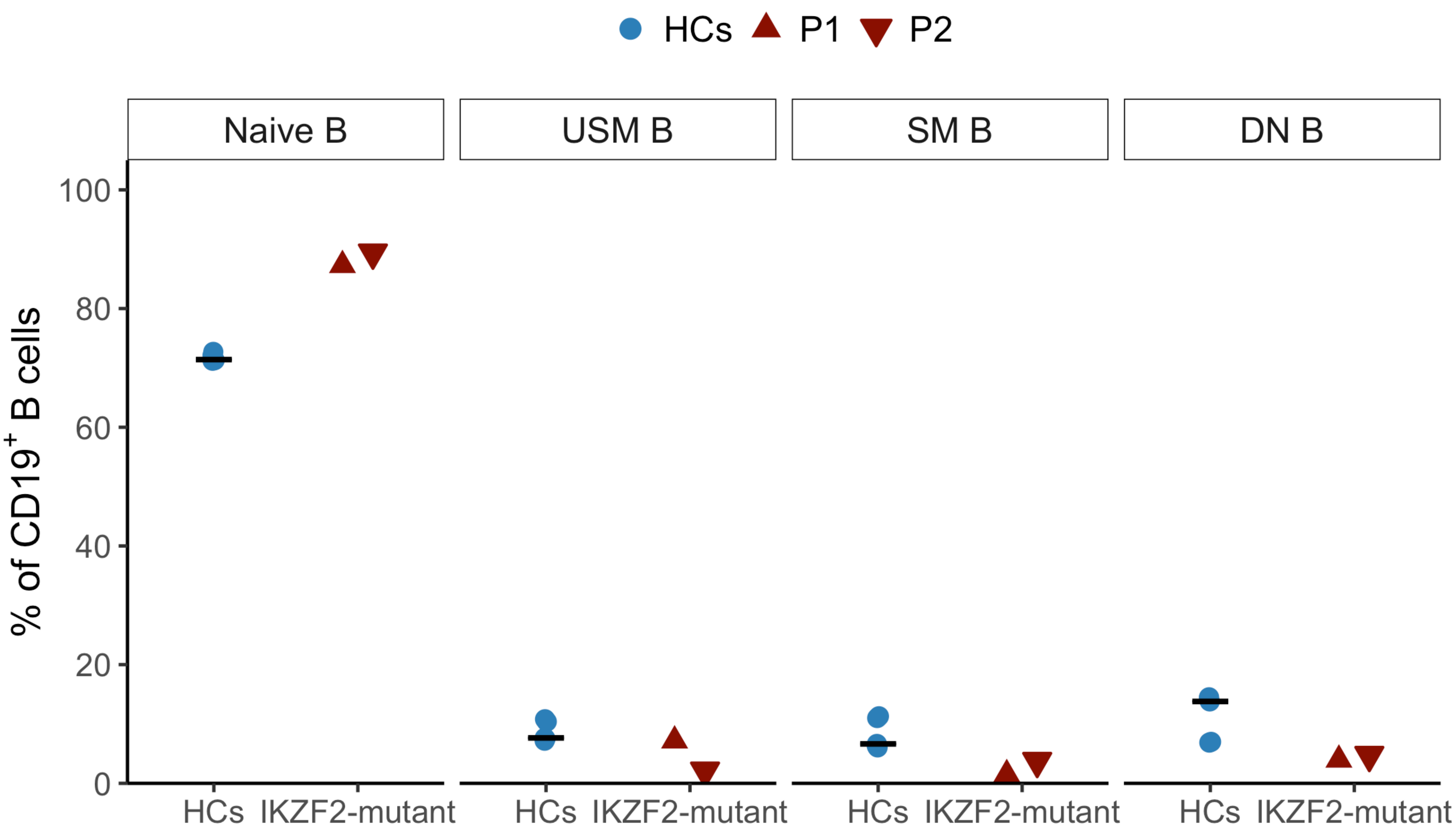
